# Disturbed mitochondrial maturation in cardiolipin remodeling-deficient cardiomyocytes

**DOI:** 10.1101/2025.08.07.668981

**Authors:** Nanami Senoo, Macie S. Sheridan, Yvonne Wohlfarter, Mackenzie T. Primrose, Emmanouil Tampakakis, Markus A. Keller, Steven M. Claypool

## Abstract

Barth syndrome, a rare X-linked genetic disorder, features early-onset cardiomyopathy. The causal gene, *TAFAZZIN*, encodes a transacylase that mediates acyl chain remodeling of cardiolipin, a critical phospholipid in the inner mitochondrial membrane. While Barth syndrome exhibits hallmark cardiolipin abnormalities, the precise mechanisms linking TAFAZZIN deficiency and disturbed cardiolipin metabolism to progressive cardiac dysfunction remain unclear. In this study, we modeled Barth syndrome cardiomyopathy in human induced pluripotent stem cell-derived cardiomyocytes with *in vitro* maturation treatments that simulate heart developmental stimuli. We found that cardiomyocyte maturation involves progressive cristae dynamics associated with protein and lipid alterations in the inner mitochondrial membrane. TAFAZZIN-deficient cardiomyocytes fail to adapt to the developmental stimuli, resulting in damaged cristae, compromised mitochondrial respiration, and cardiomyocyte dysfunction. These results demonstrate that TAFAZZIN deficiency perturbs functional and structural development of mitochondria, which may contribute to mitochondrial dysfunction and associated childhood progression to cardiomyopathy in Barth syndrome.

## Introduction

Barth syndrome (MIM#302060) is a rare, X-linked genetic disorder caused by mutations in the *TAFAZZIN* gene (Bione et al., 1996). Cardiomyopathy is the most prominent feature of Barth syndrome as it affects over 70% of patients within their first year and nearly all patients by the age of five (Clarke et al., 2013). Infancy and early childhood are critical periods, sometimes requiring cardiac transplantation (Clarke et al., 2013). Barth syndrome cardiomyopathy exhibits a spectrum of defects. While dilated cardiomyopathy is typical, hypertrophic cardiomyopathy also occurs, which can be accompanied by endocardial fibroelastosis or left ventricular noncompaction (Clarke et al., 2013; Taylor et al., 2022).

*TAFAZZIN* (TAZ) encodes a transacylase responsible for remodeling cardiolipin acyl chains. Cardiolipin, a glycerophospholipid located exclusively in the inner mitochondrial membrane (IMM), plays multifaceted, crucial roles in mitochondria, from promoting the assembly, organization, and function of the oxidative phosphorylation (OXPHOS) machinery to enabling cristae morphology (Acoba et al., 2020; Decker and Funai, 2024). Cardiolipin comprises two phosphates and four acyl chains. In Barth syndrome, TAZ deficiency disrupts cardiolipin remodeling, leading to an accumulation of monolyso-cardiolipin and a decrease in properly acylated cardiolipin. This imbalance, reflected in an elevated monolyso- cardiolipin:cardiolipin ratio, is a diagnostic hallmark of Barth syndrome (Van Werkhoven et al., 2006). In terms of acyl chains, linoleic acid dominates in normal hearts (Oemer et al., 2020); in contrast, Barth syndrome hearts display a marked decrease in linoleic acid-containing cardiolipin, instead showing other acylation patterns often with shorter and less unsaturated acyl chains (Schlame et al., 2002, 2003). These remarkable cardiolipin abnormalities are thought to perturb mitochondrial integrity, leading to destabilized and disorganized respiratory complexes and supercomplexes, and abnormal cristae morphology. However, the mechanistic link between TAZ deficiency, cardiolipin abnormalities, and Barth syndrome pathology remains to be fully elucidated.

Experimental models across organisms have been developed to study Barth syndrome, all of which feature the diagnostic elevated monolyso-cardiolipin:cardiolipin ratio (Pu, 2022). To better understand Barth syndrome pathology *in vivo*, several mouse models with targeted TAZ deletion have been developed. Doxycycline-inducible knockdown systems, while frequently used, present challenges such as incomplete TAZ depletion, variable phenotypes, and potential off-target effects (Pu, 2022). Germline TAZ knockout (KO) mice were initially reported to be lethal (Wang et al., 2020). Subsequent study has demonstrated that strain backgrounds significantly influence phenotype severity, with C57BL6/J exhibiting the most pronounced effects (Wang et al., 2023). While strain-dependent phenotypic variation mirrors the symptom spectrum in Barth syndrome patients (Clarke et al., 2013; Taylor et al., 2022), it underscores the need for careful consideration of genetic background in experimental design and data interpretation. For cardiac investigations, conditional KO strategies have been used to delete TAZ in postnatal or adult cardiomyocytes, with Myh6-Cre (Wang et al., 2020) and Xmlc2-Cre (Zhu et al., 2021) to target the Taz^flox^ allele. These conditional TAZ KO mice display cardiac dysfunction at 2-4 months of age (Wang et al., 2020; Zhu et al., 2021). A recent study established Barth syndrome patient mutation knock-in mice (TAZ G197V) (Chowdhury et al., 2023). In this model, the mutant TAZ is undetectable and cardiac dysfunction is evident at 20 weeks of age (Chowdhury et al., 2023). Strain backgrounds have not yet been considered for these conditional KO and patient mutation knock-in mice.

For investigating the molecular and cellular mechanisms underlying Barth syndrome, human induced pluripotent stem cells (hiPSCs) offer a powerful tool. Specifically, differentiating hiPSCs into cardiomyocytes allows for the study of cardiac-specific effects. Studies to date have used hiPSCs derived from Barth syndrome patients (Chowdhury et al., 2023; Dudek et al., 2016; Fatica et al., 2019; Wang et al., 2014) or engineered to delete TAZ (Sniezek Carney et al., 2024; Wang et al., 2014). Consistent with findings in patients and mouse models, these Barth syndrome model hiPSCs (Dudek et al., 2013) and differentiated cardiomyocytes (Sniezek Carney et al., 2024; Wang et al., 2014) show an elevated monolyso-cardiolipin:cardiolipin ratio. These models have revealed key cellular deficits, including impaired sarcomere integrity and contractile function (Dudek et al., 2016; Wang et al., 2014), destabilized and disorganized respiratory complexes and supercomplexes (Dudek et al., 2016; Sniezek Carney et al., 2024), and increased reactive oxygen species production (Wang et al., 2014). Alterations in mitochondrial respiration have also been reported, though with variability across studies (Chowdhury et al., 2023; Dudek et al., 2016; Sniezek Carney et al., 2024; Wang et al., 2014). The potential contribution of genetic background to TAZ-null hiPSC-derived cardiomyocyte biology has, to date, not been systematically assessed, an important omission given the broad phenotypic spectrum associated with TAZ deficiency in both outbred humans and mice (Clarke et al., 2013; Taylor et al., 2022; Wang et al., 2023).

Given the early onset of Barth syndrome cardiomyopathy, it is crucial to understand the role of TAZ in this context. To obtain detailed mechanistic insights, we challenged TAZ KO hiPSC-derived cardiomyocytes with maturation treatments that simulate the late fetal/neonatal rise of developmental stimuli. Our results using this developmental paradigm demonstrate the inability of TAZ-deficient cardiomyocytes to properly execute a stage-specific mitochondrial restructuring program, suggesting that this failure may contribute to the progressive cardiac dysfunction observed in Barth syndrome.

## Results

### Compromised mitochondrial adaptation to fatty acid exposure in TAZ-deficient cardiomyocytes

Developing hearts encounter significant changes in surrounding physiological stimuli known to promote heart maturation (Chattergoon, 2019; Girard et al., 1992; Rog-Zielinska et al., 2014). Given the early onset of Barth syndrome cardiomyopathy, we hypothesized that TAZ deficiency may disrupt cardiomyocyte adaptation to these developmental stimuli, leading to progressive dysfunction. To test our hypothesis and model this phenomenon, we obtained cardiomyocytes differentiated from wild-type (WT) and TAZ KO hiPSCs, and treated these cells with palmitic acid and L-carnitine (Pal/car) for 8 days, a treatment that mimics the fatty acid surge that occurs after birth (Girard et al., 1992; Lopaschuk and Jaswal, 2010; Zhao et al., 2019). Cardiomyocyte purity, assessed by cardiac Troponin T positive cell population, exceeded approximately 80% in both WT and TAZ KO cells, with no significant difference between genotypes (**Figure S1A**). Furthermore, cell viability was unaffected by Pal/car treatment for 8 days in both genotypes, indicating no apparent toxicity of this treatment (**Figure S1B**).

TAZ KO hiPSCs used in this study carry a 50 bp deletion at the conserved acyltransferase domain of the *TAFAZZIN* gene, as previously described (Sniezek Carney et al., 2024). Immunoblot analysis confirmed the deletion of TAZ in differentiated KO cardiomyocytes (**Figure 1A**). Additionally, we confirmed that TAZ KO cells exhibited an elevated monolyso-cardiolipin:cardiolipin ratio (**Figure 1B**) and a shift in cardiolipin molecular species towards those containing short and saturated acyl chains (**Figure S2, S3**). TAZ expression in WT cells was unchanged following Pal/car treatment for 8 days (**Figure 1A**). Consistently, the cardiolipin molecular species composition in WT and TAZ KO cells was unchanged by an 8-day Pal/car treatment (**Figure S2, S3**), although the elevated monolyso- cardiolipin:cardiolipin in TAZ KO cells was slightly normalized.

**Figure 1:**
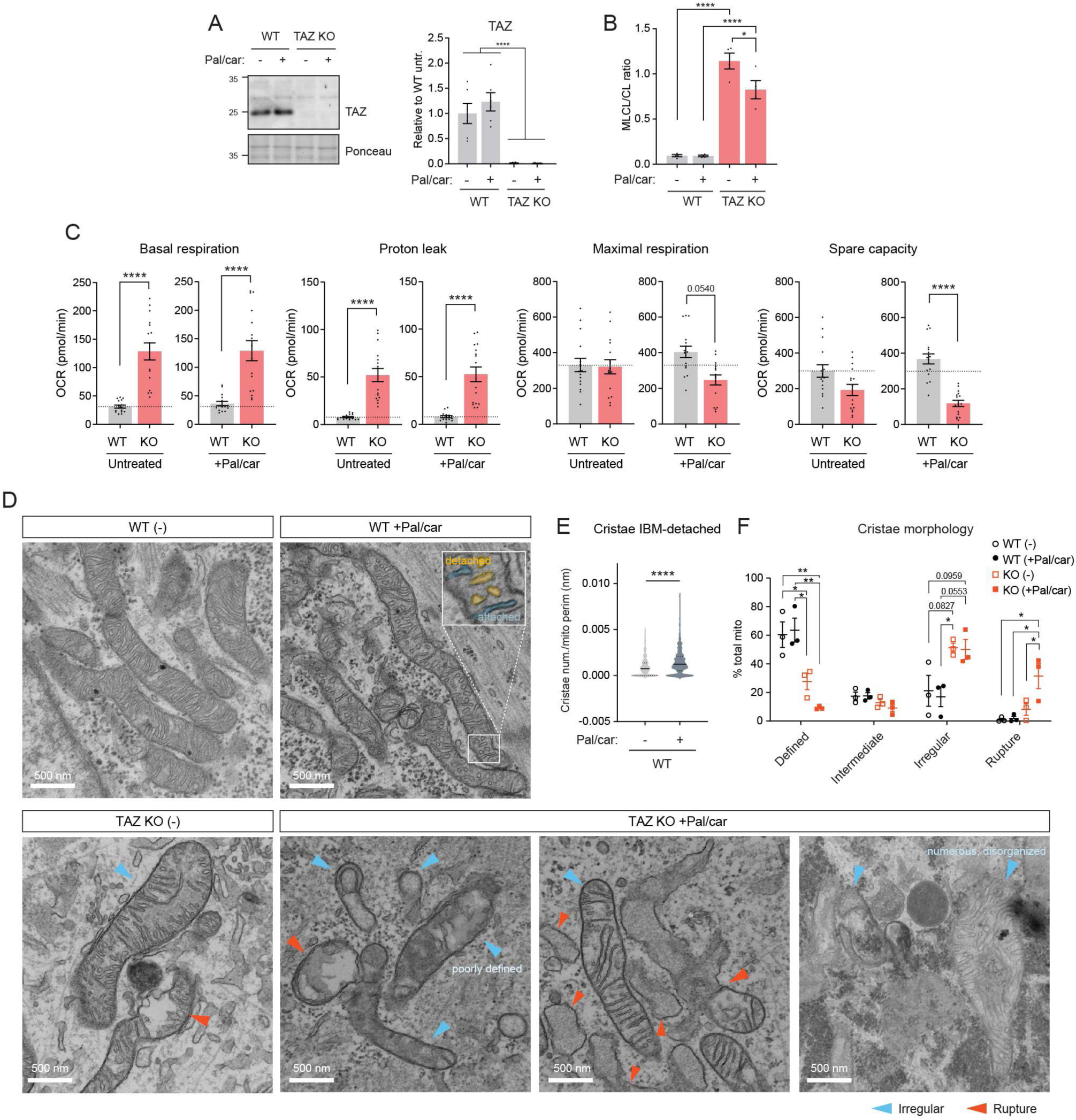
Compromised mitochondrial adaptation to fatty acid exposure in TAZ-deficient cardiomyocytes. Where indicated, WT and TAZ KO cardiomyocytes were cultured under Pal/car for 8 days. **(A)** Immunoblot analysis of TAZ expression (mean ± SEM, n = 5-6 biological replicates). **(B)** Monolyso-cardiolipin (MLCL) to cardiolipin (CL) ratio (mean ± SEM, n = 4 biological replicates). **(C)** Oxygen consumption rate (OCR) measured by Seahorse XF96e FluxAnalyzer with glucose, glutamine, pyruvate, and palmitic acid as substrates (100K/well; mean ± SEM, n = 12-16 with 4 biological replicates and 3-4 technical replicates). Dashed lines: untreated WT. **(D-F)** Cristae morphology analyzed by electron microscopy. **(D)** Representative images. See also Figure S6. **(E)** Quantification of cristae detachment from the inner boundary membrane (IBM) in WT cardiomyocytes. Data are from 3 biological replicates, with a total of 411 (untreated) and 259 (Pal/car-treated) mitochondria. Median and quartiles are shown. **(F)** Cristae morphology was categorized as follows: defined (>50% cristae with clear and aligned membranes), intermediate (25- 50% cristae with clear and aligned membranes), irregular (<25% cristae with clear and aligned membranes), and rupture (as indicated by the red arrow). Data are from 3 biological replicates, with a total of 205-442 mitochondria analyzed per sample group. Statistical significance: one-way ANOVA with Tukey’s test (A, B, C, F) and unpaired t-test (E); *p<0.05, **p<0.01, ***p < 0.001, ****p < 0.0001.

Mitochondrial respiration was measured using a Seahorse flux analyzer. Untreated TAZ KO cells showed high basal respiration and proton leak, while maximal respiration and spare capacity were comparable to WT (**Figure 1C**). Pal/car treatment to WT cells did not affect the measured parameters (**Figure 1C**). In Pal/car-treated TAZ KO cells, basal respiration and proton leak remained high; however, maximal respiration and spare capacity were significantly decreased compared to WT (**Figure 1C**). These results indicate that TAZ-deficient cardiomyocytes are recalcitrant to fatty acid exposure and lose respiration capacity in the process.

To assess substrate preference, we measured acute responses to UK-5099 (mitochondrial pyruvate carrier, MPC inhibitor) and etomoxir (carnitine palmitoyltransferase I, CPT1 inhibitor). In WT cells, Pal/car treatment caused a mild shift from pyruvate to fatty acid oxidation, as shown by increased response to etomoxir (**Figure S4A**). Immunoblot analysis of CPT and MPC showed that Pal/car treatment suppressed MPC2 expression in WT cells (**Figure S4B**), which might contribute to the shift toward fatty acid oxidation. TAZ KO cells showed a high reliance on fatty acid oxidation, which was observed under both untreated and Pal/car-treated conditions (**Figure S4A**). This suggests that substrate preference was irrelevant to the loss of respiration capacity in TAZ KO cells in response to fatty acid exposure. Although TAZ KO cells had lower levels of CPTs and MPCs (**Figure S4B**), further investigation is needed to clarify the contribution of these players to the observed respiratory phenotype. In TAZ KO cells, the steady state amounts of OXPHOS proteins, namely subunits of Complexes I, IV, and V, were decreased **(Figure S5A**), as were the respiratory supercomplexes and complex V oligomers that they help create (**Figure S5B-C**). None of these TAZ KO-associated OXPHOS expression and assembly defects were further exacerbated by Pal/car treatment (**Figure S5**). This indicates that the OXPHOS protein machinery was not involved in the altered mitochondrial respiration when TAZ KO cells were exposed to fatty acid.

Cristae morphology was analyzed using electron microscopy. In untreated WT cells, cristae were well-aligned and attached to the inner boundary membrane (IBM) (**Figure 1D**). Following Pal/car treatment to WT cells, some cristae were detached from the IBM, forming vesicular-like structures (**Figure 1D**). Quantitative analysis revealed that Pal/car treatment increased cristae detachment from the IBM in WT cells (**Figure 1E**). While TAZ KO cells showed disorganized cristae morphology even when untreated, Pal/car treatment exacerbated their morphological abnormalities (**Figure 1D**). Pal/car-treated TAZ KO cells featured a heterogeneous appearance, including poorly defined membranes, numerous disorganized membranes, and even rupture (**Figure 1D, S6**). Quantitative analysis revealed a trend for decreased ‘defined’ cristae (with clear and aligned membranes) and increased rupture (loss of compartmentalization and disruption of internal structure) in Pal/car treated TAZ KO cells (**Figure 1F**). These results indicate that fatty acid exposure stimulates cristae dynamics in cardiomyocytes that normally promotes cristae detachment. In contrast, the already abnormal cristae morphology of TAZ-deficient cardiomyocytes is exacerbated by fatty acid treatment, perhaps reflecting impaired or distorted cristae dynamics. These irregular and often ruptured mitochondria can be considered dysfunctional, likely contributing to the loss of respiration capacity in TAZ KO cells when exposed to fatty acid.

### Cardiomyocyte *in vitro* maturation increases mitochondrial respiration and cristae density

Next, we tested the impact of developmental hormones, known to rise significantly from late gestation to birth and play a crucial role in heart maturation (Chattergoon, 2019; Rog-Zielinska et al., 2014). We cultured WT and TAZ KO cardiomyocytes for 2 weeks with dexamethasone (Dex, synthetic glucocorticoid) and triiodothyronine (T3, thyroid hormone), in addition to the fatty acid exposure (Pal/car). This hormone treatment had no overt toxicity in both WT and TAZ KO cells (**Figure S7**). To validate whether this hormone treatment (Pal/car + Dex/T3 for 2 weeks) promotes cardiomyocyte maturation, we analyzed the expression of genes commonly upregulated in more mature cardiomyocytes. Pal/car + Dex/T3 treatment for 2 weeks increased TNNI3 expression (**Figure 2A, S8A**), indicating a switch in Troponin I isoform expression towards mature-type (Dark et al., 2023; Funakoshi et al., 2021; Karbassi et al., 2020). Troponin I isoform switch was also validated at the protein level; Pal/car + Dex/T3 treatment increased cardiac Troponin I (cTnI) and diminished slow skeletal muscle Troponin I (ssTnI) (**Figure 2B**). Furthermore, Pal/car + Dex/T3 treatment increased the expression of mature-type calcium and potassium handling genes (ATP2A2 and KCNJ2 respectively(Funakoshi et al., 2021; Karbassi et al., 2020)) in WT cells (**Figure 2C, S8B**). The upregulation of these genes was blunted in TAZ KO cells (**Figure 2C, S8B**), suggesting perturbation of cardiomyocyte electrophysiology that may be associated with an impact of TAZ deficiency on maturation processes. In contrast, HCN4, expressed in immature cardiomyocytes (Funakoshi et al., 2021; Karbassi et al., 2020), was low in Pal/car + Dex/T3-treated cells (**Figure 2C, S8B**). These gene expression changes were directly tied to the hormone treatment as 2 weeks of culture with Dex/T3 alone resulted in similar gene expression to that observed in Pal/car + Dex/T3-treated cells, whereas untreated or Pal/car alone-treated cells did not exhibit this response (**Figure S8**). These results demonstrate that 2-week treatment with Dex/T3 +/- Pal/car effectively promoted cardiomyocyte maturation.

**Figure 2:**
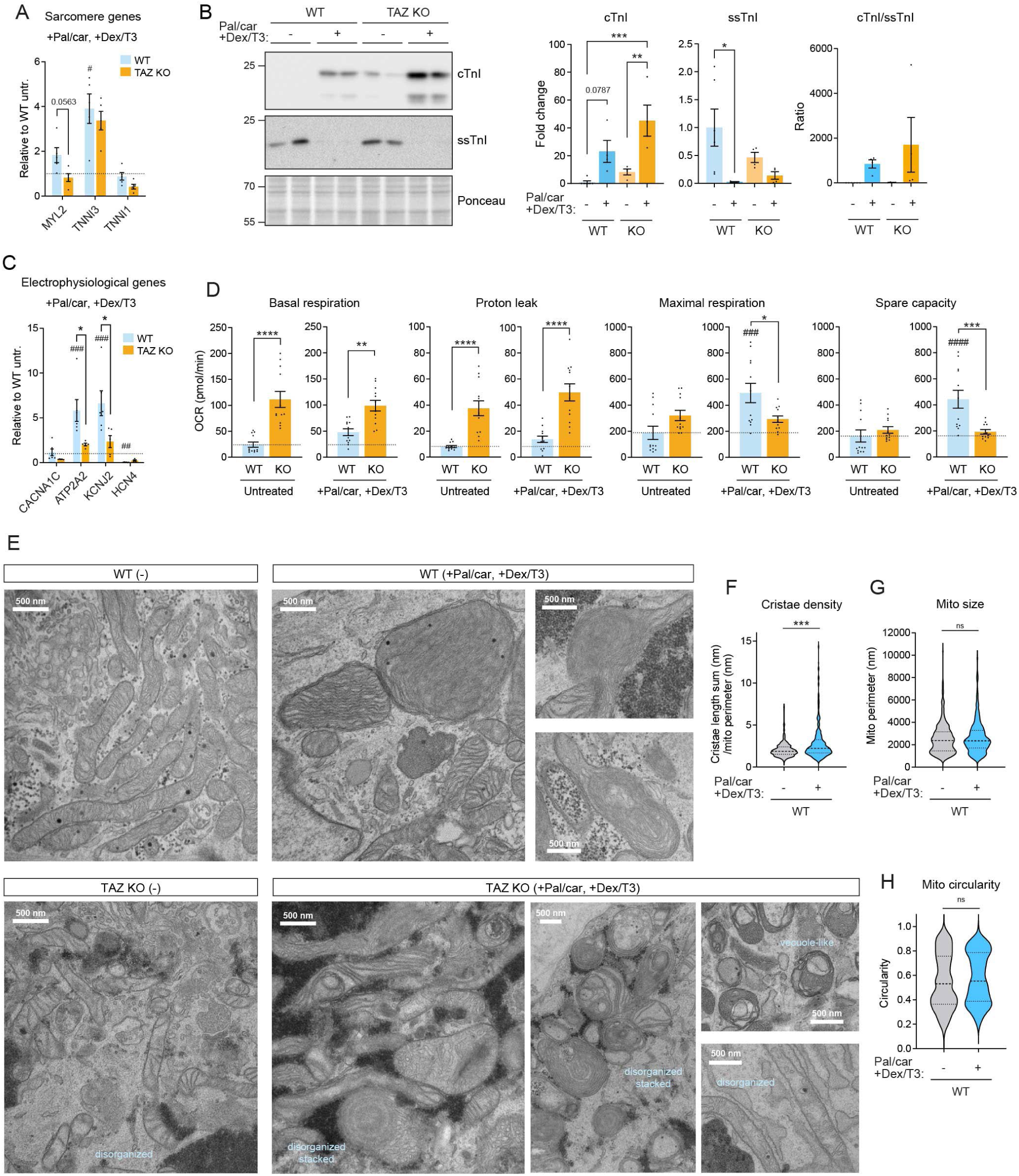
Cardiomyocyte maturation increases mitochondrial respiration and cristae density. WT and TAZ KO cardiomyocytes were cultured with Pal/car + Dex/T3 for 2 weeks as indicated. (**A**) Sarcomere gene expression (qPCR, normalized to *EEF1A1*; mean ± SEM, n = 5 biological replicates). Dashed lines: untreated WT (see Figure S8A). (**B**) Immunoblot analysis of cardiac Troponin I (cTnI) and slow skeletal muscle Troponin I (ssTnI) (mean ± SEM, n = 4-6 biological replicates). (**C**) Electrophysiological gene expression (qPCR, normalized to *EEF1A1*; mean ± SEM, n = 5 biological replicates). Dashed lines: untreated WT (see Figure S8B). (**D**) OCR measured by Seahorse XF96e FluxAnalyzer with glucose, glutamine, pyruvate, and palmitic acid as substrates (80K/well; mean ± SEM, n = 9-12 with 3 biological replicates and 3-4 technical replicates). Dashed lines: untreated WT. (**E**) Representative electron microscopy images (n = 3-4 biological replicates). (**F-H**) Quantitative analysis of mitochondrial morphology of WT cardiomyocytes. Data are from 4 biological replicates, with a total of 282 (untreated) and 260 (+Pal/car, +Dex/T3) mitochondria. Median and quartiles are shown. (**F**) Cristae density (sum of cristae length per mitochondrion). (**G**) Mitochondrial size. (**H**) Mitochondrial circularity obtained by ImageJ function (4π × area/(perimeter)^2^). Statistical significance: one-way ANOVA with Tukey’s test (A-D) and unpaired t-test (F-H); *p<0.05, **p<0.01, ***p < 0.001, ****p < 0.0001. * WT vs. TAZ KO within treatment; # vs. untreated within genotype (in A, C, D).

Having confirmed the effects of this hormone treatment on cardiomyocyte maturity, we next investigated its impact on mitochondrial respiration. In WT cells, 2-week treatment with Pal/car + Dex/T3 significantly enhanced maximal respiration and spare capacity (**Figure 2D**). In TAZ KO cells, basal respiration and proton leak remained high regardless of treatment (**Figure 2D**). Following Pal/car + Dex/T3 treatment for 2 weeks, TAZ KO cells showed lower maximal respiration and spare capacity than WT cells (**Figure 2D**); compared to untreated TAZ KO cells, these respiratory parameters were unchanged. These results demonstrate that cardiomyocyte maturation enhances mitochondrial respiration, and that TAZ deficiency hinders this process. Mitochondrial respiration of cells with Pal/car treatment alone for 2 weeks was similar to untreated cells (**Figure S9A**). Dex/T3 alone for 2 weeks resembled Pal/car + Dex/T3 treated cells (**Figure S9A**).

In addition, we assessed substrate preference following the 2-week treatments. WT cells showed high reliance on fatty acid oxidation across all treatment conditions (**Figure S9B**). TAZ KO cells also exhibited a high reliance on fatty acid oxidation, regardless of treatment (**Figure S9B**). Unlike the 8-day cultures (**Figure S4A**), the 2-week cultures induced some response to UK-5099 of TAZ KO cells, except for those treated with Pal/car alone (**Figure S9B**). This suggests that the duration of treatment may influence respiratory substrate preference in TAZ KO cells.

Cristae morphology was significantly changed following 2-week hormone treatment. This treatment produced tightly packed and layered cristae membranes in WT cells (**Figure 2E**). Quantitative analysis revealed increased cristae density in Pal/car + Dex/T3-treated WT cells, without significantly affecting mitochondrial size and circularity (**Figure 2F-H**). We noted that some mitochondria in Pal/car + Dex/T3-treated WT cells retained a similar morphology to untreated cells or displayed an apparent partial membrane densification (**Figure 2E**). This may suggest that cristae densification is ongoing, even after 2- week hormone treatment (**Figure 2E**). Strikingly, Pal/car + Dex/T3 treatment to TAZ KO cells led to marked heterogeneity in cristae morphology (**Figure 2E**). The majority of mitochondria in Pal/car + Dex/T3-treated TAZ KO cells exhibited abnormal cristae morphology, including disorganized and stacked membranes (**Figure 2E**), as well as vacuole-like compartmentalization (**Figure 2E**) that resembled structures previously reported from adult TAZ knockdown mouse hearts (Russo et al., 2024). Due to the highly disorganized and heterogeneous nature of cristae, detailed quantitative cristae morphology analysis for TAZ KO cells was not feasible. These observations indicate that Pal/car + Dex/T3 treatment to promote cardiomyocyte maturation leads to cristae densification, which is severely disrupted in TAZ deficiency.

### Cardiomyocyte *in vitro* maturation develops the inner mitochondrial membrane protein machinery

IMM is rich in protein machinery, with the five OXPHOS complexes and adenine nucleotide translocator (ANT) constituting approximately two-thirds of all IMM proteins in bovine heart (Schlame, 2021). We observed a differential impact of the 2-week hormone treatment on IMM protein machinery in WT and TAZ KO cells. Pal/car + Dex/T3 treatment to WT cells increased steady-state levels of Complexes I and II subunits (**Figure 3A, S10A**) and Complex IV (**Figure 3B-C**). At baseline, TAZ KO cells exhibited significantly lower steady-state levels of Complexes I, IV, and V subunits compared to WT cells (**Figure 3A, S10A**). However, Pal/car + Dex/T3 treatment to TAZ KO cells increased the level of Complex II subunit and showed a trend toward normalizing the levels of Complexes I, III, and V subunits (**Figure 3A, S10A**). Blue native-PAGE analysis of individual complexes further supported this finding, demonstrating that the abundances of Complexes I, III, and IV were normalized by Pal/car + Dex/T3 treatment in TAZ KO cells (**Figure 3B-C**).

**Figure 3:**
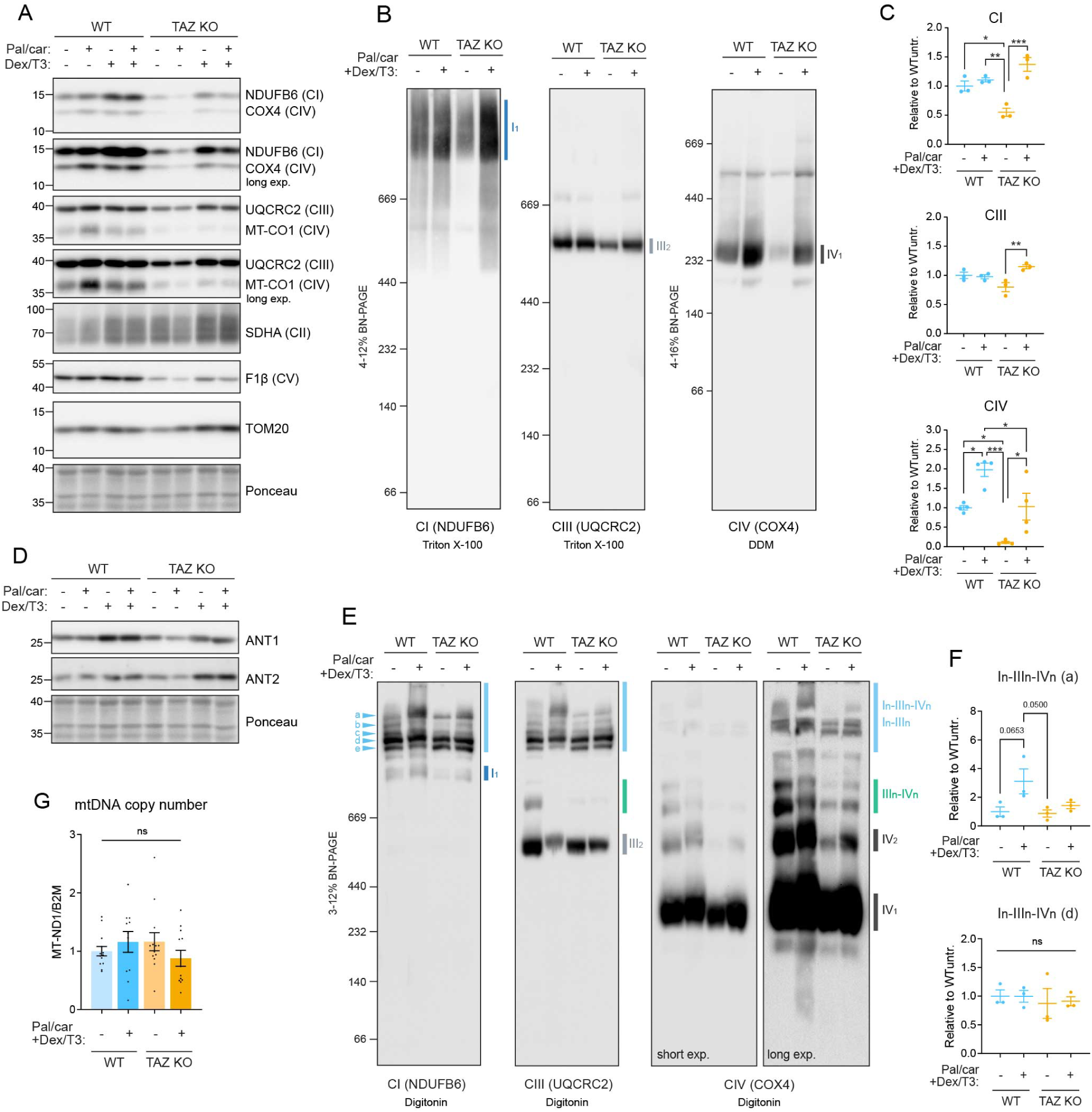
Cardiomyocyte maturation develops the inner mitochondrial membrane protein machinery. WT and TAZ KO cardiomyocytes were cultured with the indicated treatments for 2 weeks. **(A)** Immunoblot analysis of OXPHOS subunits (Complexes I, II, III, IV, and V). Representative images from n = 5-6 biological replicates are shown. **(B-C)** Individual respiratory complexes. Triton X-100 or n-dodecyl-β-D-maltoside (DDM)-solubilized cell extracts (protein:detergent = 1:4) were analyzed by blue native-PAGE and immunoblot. **(D)** Immunoblot analysis of adenine nucleotide translocators. Representative images from n = 5-6 biological replicates are shown. **(E-F)** Respiratory supercomplex assembly. Digitonin-solubilized cell extracts (protein:digitonin = 1:4) were analyzed by blue native-PAGE and immunoblot. Numerical subscripts specify the number of individual complex units comprising the indicated assembly; ’n’ denotes a variable number. Quantifications were performed on immunoblots against NDUFB6 (mean ± SEM, n = 3 biological replicates). **(G)** Mitochondrial DNA copy number analyzed by qPCR (mean ± SEM, n = 3-5 biological replicates). Statistical significance: one-way ANOVA with Tukey’s test; *p<0.05, **p<0.01, ***p < 0.001.

The hormone treatment also differentially affected the abundance of ANT isoforms, a carrier that exchanges ADP and ATP across the IMM. We assessed ANT1 and ANT2, the two major isoforms among four in humans. ANT1 is predominantly expressed in heart and skeletal muscle, while ANT2 is ubiquitously expressed (**Figure S11**) (Cardoso-Moreira et al., 2019; Graham et al., 1997; Stepien et al., 1992). Pal/car + Dex/T3 treatment increased ANT1 levels in both WT and TAZ KO cells, but induced ANT2 levels only in TAZ KO cells (**Figure 3D, S10B**).

Furthermore, we analyzed respiratory supercomplex assembly by blue native-PAGE. Pal/car + Dex/T3-treated WT cells showed an abundant high-order supercomplex, composed of Complexes I, III, and IV (indicated as In-IIIn-IVn (a) in **Figure 3E-F**). This appeared to be facilitated by an altered Complex III population, as the abundance of smaller supercomplexes (IIIn-IVn) and Complex III homodimer (III2) were diminished in Pal/car + Dex/T3-treated WT cells. The high-order supercomplex was barely induced in TAZ KO cells by Pal/car + Dex/T3 treatment, with a substantial amount of Complex III homodimer (III2) remaining (**Figure 3E-F**). This observation was particularly noteworthy: despite the induction of individual complex abundances **(Figure 3C**), these complexes failed to assemble into the supercomplex in the absence of TAZ. Complex V assembly was not affected by the hormone treatment in WT cells. Untreated TAZ KO cells showed disorganized Complex V assembly, which was partially restored by the hormone treatment (**Figure S10C**).

These findings suggest that cardiomyocyte maturation promotes enrichment and assembly of IMM proteins, and that TAZ deficiency perturbs this process. Note that the effect of Pal/car + Dex/T3 treatment on overall mitochondrial content in WT cells was minimal. The abundance of TOM20, a protein in the outer mitochondrial membrane (OMM), did not change in WT cells across treatments (**Figure 3A, S5A, S10A**). In TAZ KO cells, TOM20 abundance was comparable to WT with the hormone treatments (**Figure 3A, S5A, S10A**). Mitochondrial DNA copy number remained consistent across genotypes and treatments (**Figure 3G, S10D**), indicating that changes in IMM protein machinery were not due to changes in mitochondrial number.

### Cardiomyocyte *in vitro* maturation drives cardiolipin remodeling

To elucidate the influence of cardiomyocyte maturation on cardiolipin, we conducted a detailed analysis of cardiolipin and monolyso-cardiolipin species. Treatment with Pal/car + Dex/T3 significantly altered cardiolipin molecular species in WT cells, evidenced by the distinct separation observed in principal component analysis (**Figure 4A**). Specifically, Pal/car + Dex/T3-treated WT cells exhibited a transition from cardiolipin species rich in unsaturated acyl chains to those with shorter and less unsaturated acyl chains (**Figure 4A, S12**). TAZ KO cells retained a high, albeit slightly reduced, monolyso- cardiolipin:cardiolipin ratio following Pal/car + Dex/T3 treatment (**Figure 4B**). The impact of this treatment on the overall cardiolipin composition in TAZ KO cells was minimal (**Figure 4A, S12**). Immunoblot analysis revealed an increase in TAZ expression in hormone-treated WT cells (**Figure 4C**) that was in part, transcriptionally mediated (**Figure 4D**). These results suggest that the hormone treatment induces cardiolipin remodeling in concert with cardiomyocyte maturation, with TAZ playing a critical role in this process. Indeed, the hormone treatment upregulated the expression of genes involved in cardiolipin biosynthesis and remodeling (Acoba et al., 2020; Decker and Funai, 2024; Ren et al., 2025) in WT cells (**Figure 4D**). Future research should characterize the contributions of pathways beyond TAZ in this context.

**Figure 4:**
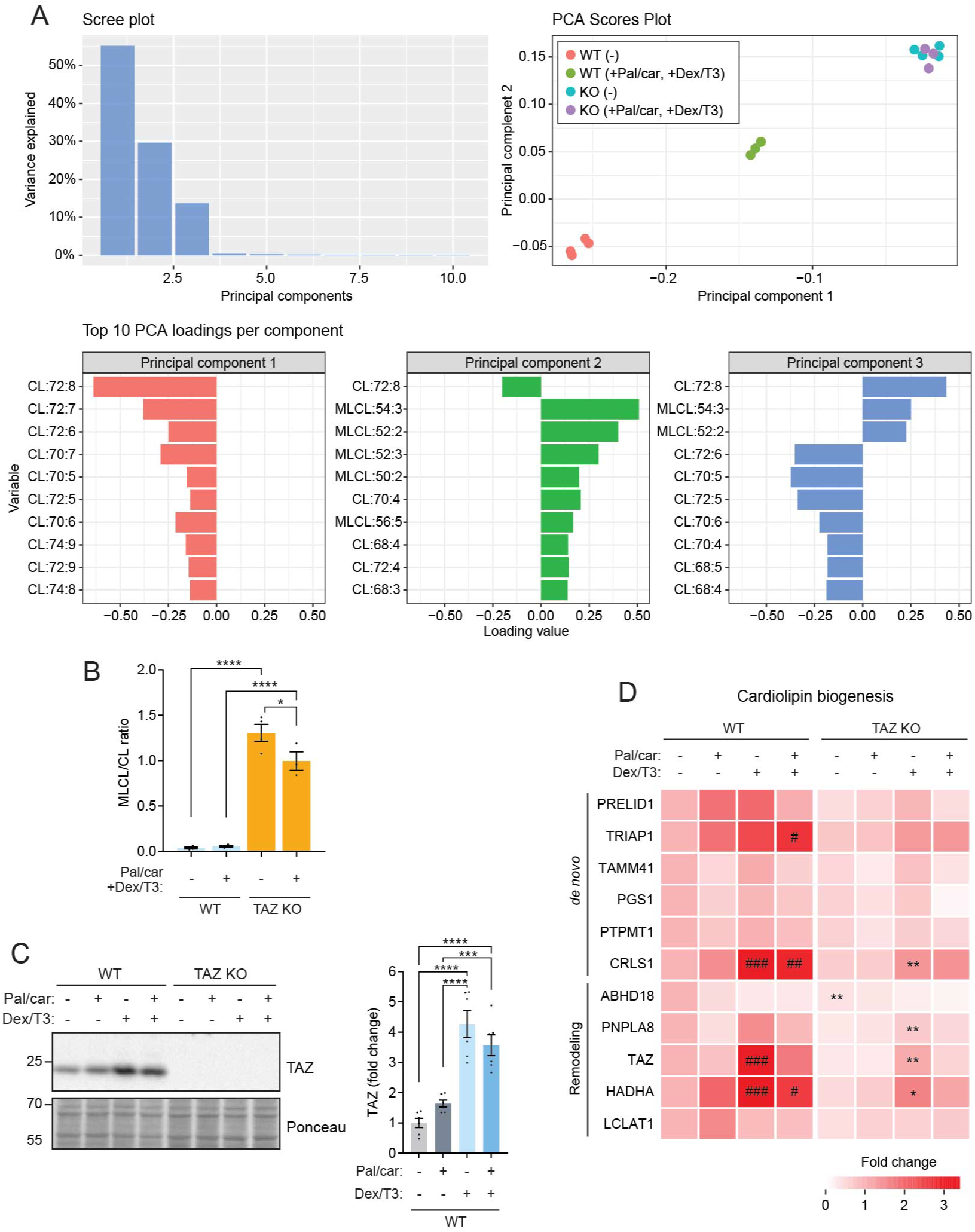
Cardiomyocyte maturation drives cardiolipin remodeling. WT and TAZ KO cardiomyocytes were cultured for 2 weeks under indicated conditions. **(A)** Principal component analysis (PCA) of cardiolipin and monolyso-cardiolipin species. **(B)** Monolyso-cardiolipin (MLCL) to cardiolipin (CL) ratio (mean ± SEM, n = 3-4 biological replicates). **(C)** Immunoblot analysis of TAZ (mean ± SEM, n = 6 biological replicates). **(D)** Heatmap of cardiolipin biogenesis gene expression (qPCR, normalized to *EEF1A1*; mean, n=3-6 biological replicates). Statistical significance: one-way ANOVA with Tukey’s test; *p<0.05, **p<0.01, ***p < 0.001, ****p < 0.0001. * WT vs. TAZ KO within treatment; # vs. untreated within genotype (in D).

To further clarify the role of TAZ in mitochondrial development associated with cardiomyocyte maturation, we re-expressed WT or mutant TAZ (H69Q) in TAZ KO hiPSCs and differentiated them into cardiomyocytes. TAZ H69Q is a catalytically inactive mutation tied to Barth syndrome (Lu et al., 2016). We obtained two WT TAZ re-expression clones (clones 1 and 3) and one H69Q clone. Upon the 2-week Pal/car + Dex/T3 treatment, WT TAZ rescue clones were expressed lower than endogenous TAZ in WT cells, while TAZ H69Q expression was comparable to endogenous TAZ levels (**Figure 5A**). Compared to TAZ KO cells, the monolyso-cardiolipin:cardiolipin ratio was significantly decreased in WT TAZ re-expressing cells, but remained high in H69Q mutant cells (**Figure 5B**). High basal respiration and proton leak observed in TAZ KO cells were reversed by re-expressing WT TAZ but not the H69Q mutant (**Figure 5C**). Re-expressing WT TAZ restored the spare capacity of TAZ KO cells, while the H69Q mutant did not (**Figure 5C**). These results formally demonstrate that TAZ is essential for proper mitochondrial respiration and that this function depends on its catalytic activity.

**Figure 5:**
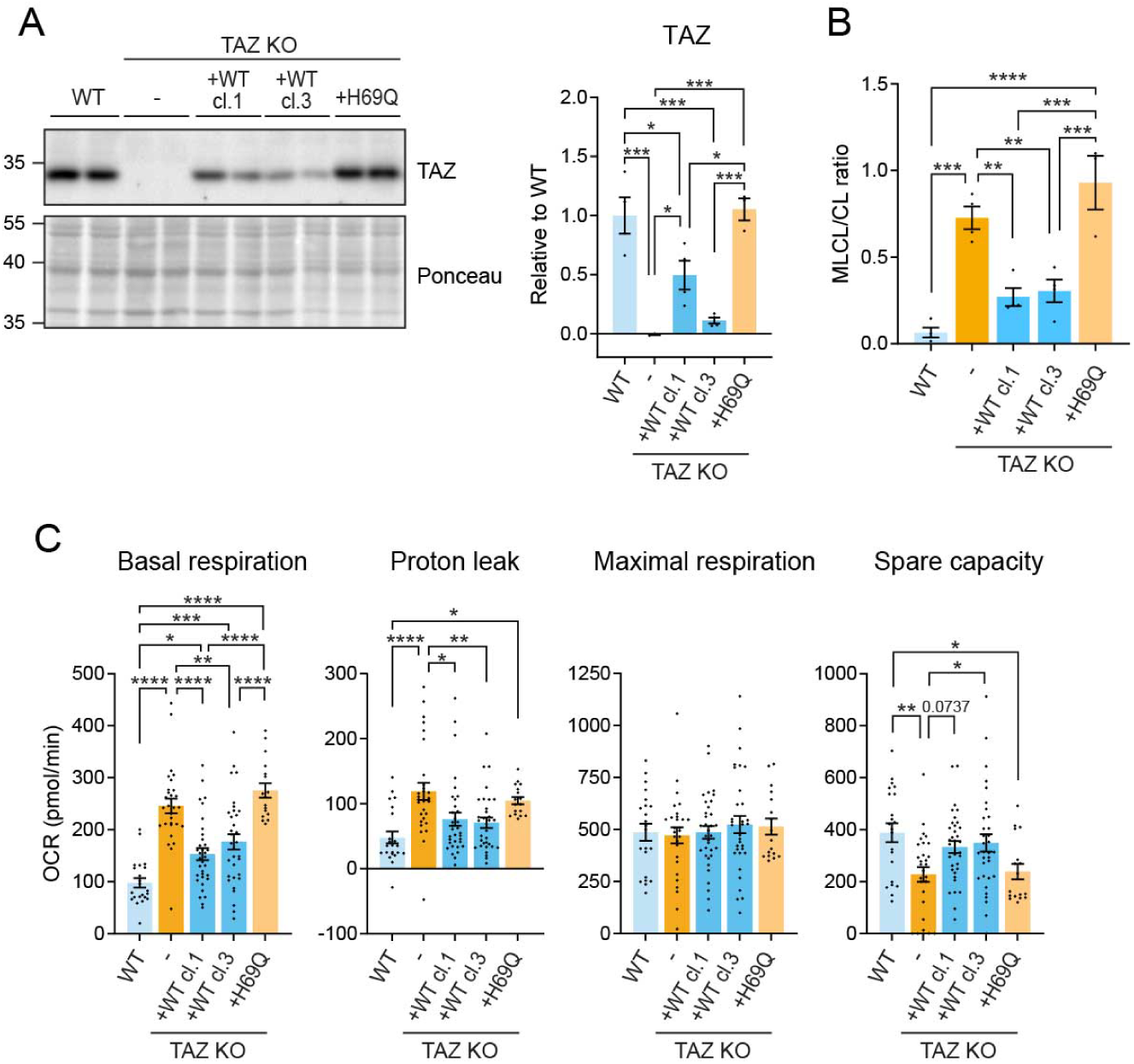
TAZ-mediated cardiolipin remodeling is essential for proper mitochondrial respiration. The indicated iPSC lines were differentiated into cardiomyocytes and treated with Pal/car + Dex/T3 for 2 weeks. **(A)** Immunoblot analysis of TAZ (mean ± SEM, n = 3-4 biological replicates). **(B)** Monolyso-cardiolipin (MLCL) to cardiolipin (CL) ratio (mean ± SEM, n = 3-4 biological replicates). **(C)** OCR measured by Seahorse XF96e FluxAnalyzer with glucose, glutamine, pyruvate, and palmitic acid as substrates (80K/well; mean ± SEM, n = 17-34 with 3-5 biological replicates and 2-6 technical replicates). Statistical significance: one-way ANOVA with Tukey’s test; *p<0.05, **p<0.01, ***p < 0.001, ****p < 0.0001.

### TAZ deficiency causes cardiomyocyte dysfunction

Next, we evaluated functional consequences of TAZ-deficient cardiomyocytes as a function of maturation. Gene expression analysis of natriuretic peptides (NPPA and NPPB), markers of heart stress and failure, showed no difference between WT and TAZ KO cells at 8 days of culture (**Figure 6A**). Following 2 weeks, NPPA and NPPB expression was markedly higher in TAZ KO cells, especially those treated with Pal/car+ Dex/T3 (**Figure 6B**). This trend was also observed in untreated TAZ KO cells (**Figure 6B**), suggesting a potential influence of long-term culture on hiPSC-derived cardiomyocyte maturity, as previously reported (Venkatesh et al., 2021). After 2 weeks of culture with Pal/car + Dex/T3, TAZ KO cells showed significantly elevated proBNP secretion (**Figure 6C**). This elevation was reversed by re-expressing WT TAZ (**Figure 6C**). Additionally, we cultured the cells under high energetic demand by treating them with endothelin-1 (ET-1). ET-1, a vasoconstrictor peptide, induces hypertrophic responses in hiPSC-derived cardiomyocytes, resulting in diastolic dysfunction (Carlson et al., 2013; Johansson et al., 2020; Redwanz et al., 2024). Under ET-1, all genotypes showed comparably high proBNP secretion (**Figure 6C**).

**Figure 6:**
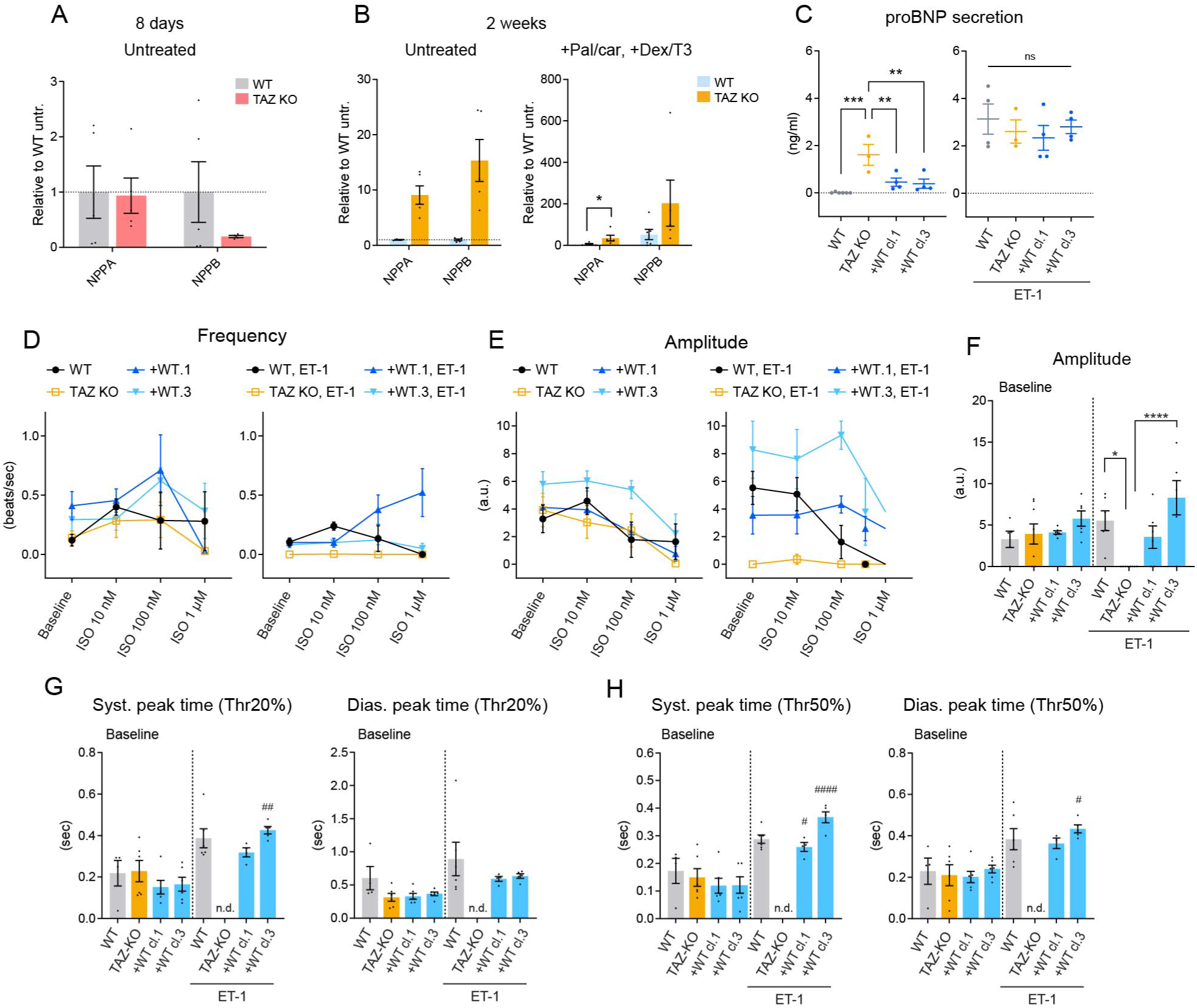
TAZ deficiency causes cardiomyocyte dysfunction. **(A)** *NPPA* and *NPPB* gene expression (qPCR, normalized to *YWHAZ*) in WT and TAZ KO cardiomyocytes cultured for 8 days (mean ± SEM, n = 5 biological replicates). **(B)** *NPPA* and *NPPB* gene expression (qPCR, normalized to *EEF1A1*) after 2 weeks of treatment (mean ± SEM, n = 5 biological replicates). Dashed lines: untreated WT. **(C-H)** Cardiomyocytes derived from the indicated iPSC strains were cultured under Pal/car + Dex/T3 for 2 weeks with or without 10 nM endothelin-1 (ET-1). (**C**) Secreted proBNP levels in the culture supernatant were quantified by ELISA (mean ± SEM, n = 3-6 biological replicates). (**D-H**) Cardiomyocyte contractions were analyzed by video recordings. Monolayered cardiomyocyte sheets were stimulated with increasing concentrations of isoproterenol (ISO) (mean ± SEM, n = 5-8 biological replicates). (**D**) Contraction frequency. (**E**) Contraction amplitude. (**F**) Amplitude at baseline. (**G, H**) Systolic and diastolic peak times at 20% (G) and 50% (H) thresholds at baseline. Limited response in ET- 1 treated TAZ KO cells precluded data display (n.d., not detected). Statistical significance: one-way ANOVA with Tukey’s test; *p<0.05, **p<0.01, ***p < 0.001, ****p < 0.0001. * Genotype comparison within treatment; # ± ET-1 comparison within genotype (in F-H).

To investigate cardiomyocyte contractile ability, we video-recorded spontaneous contractions of monolayered cardiomyocyte sheets cultured under Pal/car + Dex/T3 for 2 weeks. In addition to baseline assessment, isoproterenol (ISO), a beta-adrenergic receptor agonist, was used to stimulate contractions. At baseline, WT and TAZ KO cells contracted at ∼ 0.1 beats/sec (**Figure 6D-E**), while TAZ KO cells displayed significant replicate-to-replicate variability (**Figure S13A**). WT cells responded to 10 nM ISO with increased contraction frequency, while higher ISO concentrations diminished contractility (**Figure 6D-E, S13A**). TAZ KO cells showed replicate-to-replicate variability in their responses to ISO (**Figure S13A**), similar to baseline contraction. This irregular and heterogeneous behavior, which resembles behaviors observed in hiPSC-derived cardiomyocytes generated from patients with cardiac dysfunction (Mekies et al., 2021; Schick et al., 2018; Word et al., 2021) suggests TAZ KO cells have compromised contractility. Re-expression of WT TAZ in TAZ KO cells restored the response to ISO stimulation, although the most pronounced effect was observed at 100 nM, a higher concentration than what optimally induced WT cell contraction (**Figure 6D-E**). WT cells treated with ET-1 showed a slightly blunted response to ISO; however, ET-1 treatment of TAZ KO cells nearly abolished contractility (**Figure 6D-E, S13B**). This suggests that TAZ KO cells are susceptible to stress under high energetic demand. Further analysis of contraction peaks showed no differences in amplitude or peak times between genotypes in the absence of ET-1 (**Figure 6F-H**). Treated with ET-1, WT TAZ rescued cells showed elongated systolic or diastolic peak times (**Figure 6G-H**). Due to their abolished contractility, we failed to quantitatively assess contraction peaks of ET-1-treated TAZ KO cells. Collectively, the results demonstrate that TAZ KO cells exhibit impaired cardiomyocyte function upon maturation.

### Genetic background modulates the phenotypic spectrum of TAZ-deficient cardiomyocytes

There is a broad phenotypic spectrum associated with TAZ deficiency in outbred humans and mice (Clarke et al., 2013; Taylor et al., 2022; Wang et al., 2023). In this context, we asked whether the TAZ KO cardiomyocyte phenotypes documented above were also evident in another parental hiPSC line (C11) genetically engineered to lack *TAFAZZIN* using the same strategy as (Sniezek Carney et al., 2024). Following the same methodologies, we differentiated C11 WT and two TAZ KO hiPSC clones into cardiomyocytes and employed the 2-week hormone maturation treatment (Pal/car + Dex/T3). As expected, TAZ protein was not detected in either TAZ KO cells which importantly, also displayed an elevated monolyso-cardiolipin:cardiolipin ratio that is diagnostic of TAZ deficiency (**Figure 7A, B**). C11 WT and TAZ KO cells exhibited comparable cardiomyocyte purity, as indicated by cardiac Troponin T positive cell population exceeding 80% (**Figure S14A**). Cell viability analysis showed that C11 cells were slightly sensitive to the hormone maturation treatment (**Figure S14B**). C11 WT cells showed increased TAZ expression following hormone treatment (**Figure 7A**), consistent with the results observed in the previous cell line (**Figure 4C**).

**Figure 7:**
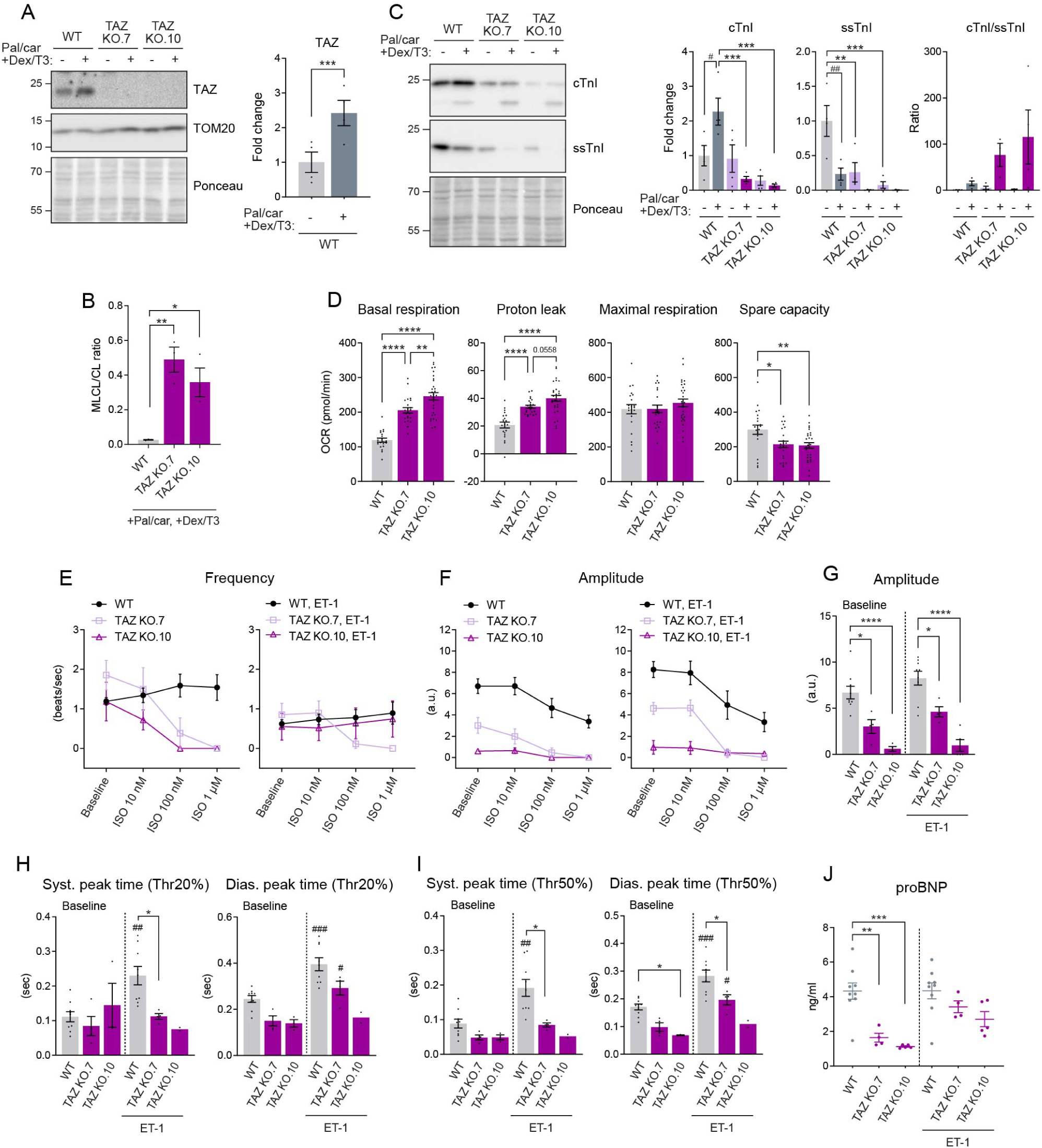
Genetic background modulates the phenotypic spectrum of TAZ-deficient cardiomyocytes. C11 WT and TAZ KO cardiomyocytes were cultured under Pal/car +Dex/T3 for 2 weeks. **(A)** Immunoblot analysis of TAZ (mean ± SEM, n = 4 biological replicates). **(B)** Monolyso-cardiolipin (MLCL) to cardiolipin (CL) ratio (mean ± SEM, n = 3 biological replicates). **(C)** Immunoblot analysis of cardiac Troponin I (cTnI) and slow skeletal muscle Troponin I (ssTnI) (mean ± SEM, n = 4 biological replicates). **(D)** OCR measured by Seahorse XF96e FluxAnalyzer with glucose, glutamine, pyruvate, and palmitic acid as substrates (40K/well; mean ± SEM, n = 21-29 with 4-5 biological replicates and 5-6 technical replicates). **(E-I)** Indicated cardiomyocytes were cultured under Pal/car + Dex/T3 with or without 10 nM ET-1. Cardiomyocyte contractions were analyzed by video recordings with ISO stimulation (mean ± SEM, n = 4-9 biological replicates). **(E)** Contraction frequency. (**F**) Contraction amplitude. **(G)** Baseline amplitude. **(H, I)** Systolic and diastolic peak times at 20% (H) and 50% (I) thresholds at baseline. Limited response in ET-1 treated TAZ KO.10 cells (n=2/5) precluded error bar and statistical significance display. **(J)** Secreted proBNP levels in the culture supernatant were quantified by ELISA (mean ± SEM, n = 4-9 biological replicates). Statistical significance: one-way ANOVA with Tukey’s test; *p<0.05, **p<0.01, ***p < 0.001, ****p < 0.0001. * Genotype comparison within treatment; # ± ET-1 comparison within genotype (in G-J).

To evaluate cardiomyocyte maturity, we analyzed Troponin I isoform expression (**Figure 7C**). C11 WT cells treated with Pal/car + Dex/T3 showed the expected maturation-related shift from ssTnI to cTnI, indicating enhanced cardiomyocyte maturity. Following Pal/car + Dex/T3 treatment, ssTnI expression, which was lower in TAZ KO than WT at baseline, was reduced in both genotypes. In contrast, cTnI expression was increased in WT cells, as expected, but not in TAZ KO cells in response to 2-week treatment. The distorted expression of Troponin I isoforms in C11 TAZ KO cells suggests potential dysregulation of sarcomere integrity (Crocini and Gotthardt, 2021) and/or inefficient maturation.

We measured mitochondrial respiration of C11 WT and TAZ KO cells after the 2-week hormone treatment. C11 TAZ KO cells showed high basal respiration, high proton leak, and low spare capacity (**Figure 7D**), consistent with results obtained from the previous cell line (**Figure 2D, 5C**).

The contractile behavior of C11 cells differed from that observed in the previous cell line. At baseline, C11 WT cells exhibited frequent contractions at ∼1.1 beats/sec (**Figure 7E, S15**). C11 TAZ KO cells also contracted and their beating pattern was relatively consistent across replicates (**Figure 7E-F, S12**), unlike the more variable pattern observed in the previous parental TAZ KO cells (**Figure 6D-E, S11**). Notably, the beat amplitude of C11 TAZ KO cells was lower than that of WT cells (**Figure 7F-G**). ISO stimulation at a low concentration (10 nM) had only a minimal impact on the contractility of WT and TAZ KO cells (**Figure 7E-F**). Higher concentrations of ISO (>100 nM) further increased the beat frequency of C11 WT cells while decreasing their amplitude (**Figure 7E-F**). In contrast, the same higher concentrations of ISO markedly decreased the beat frequency of C11 TAZ KO cells (**Figure 7E-F**), suggesting a decreased ability of these cells to meet high energy demands. Following ET-1 treatment, C11 WT cells exhibited elongated systolic and diastolic peak times (**Figure 7H-I**). Although C11 TAZ KO cells continued to contract under ET-1 treatment, their contraction amplitude was lower than that of WT cells (**Figure 7E-G**). Interestingly, the proBNP secretion pattern in the C11 cell line differed from that observed in the previous cell line. While C11 WT cells secreted high levels of proBNP, TAZ KO cells secreted lower levels (**Figure 7J**). ET-1 treatment did not affect proBNP secretion in either genotype. We speculate that the parental-dependent phenotypic variation captured here may partially reflect the spectrum of Barth syndrome patients’ symptoms.

## Discussion

In this study, we modeled Barth syndrome cardiomyopathy using hiPSC-derived cardiomyocytes to explore the disease mechanisms underlying early-onset and progressive cardiac dysfunction. Maturation treatments simulating heart developmental stimuli revealed staged developmental adaptations of mitochondria in cardiomyocytes, involving progressive cristae dynamics. Exposure to fatty acid caused cristae isolation from the IBM, while hormone treatment led to cristae densification along with enhanced cardiomyocyte maturity. In contrast, maturation treatments to TAZ-deficient cardiomyocytes resulted in highly irregular and damaged cristae morphology, indicating a perturbation of cristae dynamics. Following maturation treatments, TAZ-deficient cardiomyocytes showed impaired mitochondrial respiration capacity and compromised contractile ability. Our findings indicate that TAZ plays a crucial role in the proper mitochondrial development that occurs as cardiomyocytes mature – the inability of TAZ-deficient cardiomyocytes to robustly undergo these processes may underlie progressive cardiac dysfunction in Barth syndrome.

Our results highlight the significance of cristae dynamics in cardiomyocyte maturation. Step-wise cristae dynamics may allow for cardiomyocytes i) to metabolically adapt to physiological stimuli that fluctuate widely in early developmental stages; and ii) to achieve the high energetic capacity characteristic of the heart as it further develops. The concept of cristae dynamics emerged in the 1960s through electron microscopy studies and has been further elaborated by recent advancements in live-cell imaging with super-resolution nanoscopy (Hackenbrock, 1966; Mannella, 2006; Kondadi et al., 2020a). Recent studies have shown that cristae undergo fission and fusion and can even exist as isolated vesicles, and that cristae dynamics are influenced by cellular energetic status (Golombek et al., 2024; Kondadi et al., 2020b; Wolf et al., 2019). Although we observed that fatty acid exposure increased cristae isolation in cardiomyocytes (**Figure 1D-E**), the physiological significance of cristae isolation remains unclear. As discussed previously (Kondadi et al., 2020a; Reichert, 2024), potential benefits include enhanced proton trapping for ATP synthesis. Switching between connected and isolated cristae states may be repeated under fluctuating energy demands, as permanent isolation would limit the exchange of metabolites between the mitochondrial matrix and cytosol. Presumably, fatty acid-stimulated cristae isolation in cardiomyocytes represents a transient adaptive response to fatty acid surge that occurs postnatally. Aligned with this idea, isolated, vesicle-like cristae were observed in developing mouse hearts, particularly a few days after birth (Neary et al., 2014). Upon maturation, we found cristae densification in cardiomyocytes (**Figure 2**). Dense cristae are characteristic of healthy mature hearts (Hoppel et al., 2009); in contrast, cristae damage (a decrease in cristae density) has been associated with heart diseases (Collins et al., 2021) and aging (Hinton et al., 2023). Previous reports have shown increased cristae density upon maturation in hiPSC-derived cardiomyocytes (Funakoshi et al., 2021) and during development in mouse and rabbit hearts (Zhao et al., 2019; Neary et al., 2014; Smith and Page, 1977). Concomitantly with cristae densification, our mature cardiomyocytes exhibited enhanced mitochondrial respiration, as well as IMM protein enrichment and respiratory supercomplex formation (**Figure 3**). This suggests that cardiomyocyte maturation-associated structural, organizational, and functional modifications to mitochondria ‒ which we collectively refer to as cardiac mitomorphosis ‒ are needed to optimize energetic capacity to meet the relentless demands of the heart.

Irregular and damaged cristae morphology observed in our TAZ-deficient cardiomyocytes may reflect a perturbation of cristae dynamics, emphasizing a critical role of TAZ for this membrane-contorting capacity. Our observations are generally consistent with those in Barth syndrome patients and animal models alike. Cristae abnormalities reported in the previous studies are diverse, likely affected by differences in developmental stages and cardiomyopathy progression status. Barth syndrome patients with severe cardiomyopathy exhibit a spectrum of cristae abnormalities, including reduced density, compartmentalization, lamellar or stacked structures, and giant mitochondria (Bissler et al., 2002; Clarke et al., 2013). Similarly, mouse TAZ KO hearts display irregular cristae structures, including onion-shaped formations (Zhu et al., 2021; Wang et al., 2020). TAZ knockdown resulted in less dense cristae and vacuole-like compartmentalization (Russo et al., 2024).

Cardiolipin biogenesis and cristae structure are intricately linked. As previously demonstrated, the intramitochondrial phospholipid transport pathway crucial for delivering cardiolipin precursors to the IMM, contributes to cristae formation; conversely, depletion of cristae structure leads to a reduction in cardiolipin levels (Kojima et al., 2019). Aligning with this paradigm, we found a remodeling of cardiolipin molecular species in WT cells as they undergo maturation (**Figure 4**). A reciprocal mechanism likely underlies this change: cristae densification during cardiomyocyte maturation establishes an environment conducive to cardiolipin remodeling, which, in turn, facilitates cristae formation. Our data indicate that TAZ drives this maturation-associated cardiolipin alteration, in substantial accordance with a prior study (Xu et al., 2021). Moreover, the *de novo* cardiolipin synthesis pathway, including intramitochondrial phospholipid transport and the terminal step pre-acyl chain remodeling, likely participates in this process (**Figure 4D**), though further characterization is required to fully elucidate their involvement. Interestingly, while cardiolipin in normal hearts is predominantly composed of linoleic acid (Oemer et al., 2020), the maturation-associated cardiolipin remodeling under Pal/car + Dex/T3 treatment yielded shorter and less unsaturated cardiolipins (**Figure 4A, S12**). This observation suggests that maturation-driven cardiolipin remodeling preferentially incorporates fatty acids that are abundant in the cellular environment, such as palmitic acid and its derivatives featured in our system. This aligns with a previously proposed model highlighting the influence of the cellular lipid environment on cardiolipin molecular species composition (Oemer et al., 2020).

Our TAZ KO hiPSC-derived cardiomyocytes exhibited an elevated monolyso- cardiolipin:cardiolipin ratio, a hallmark of Barth syndrome as documented in patient samples and other experimental models (Pu, 2022). Previous studies using TAZ-deficient heart models have shown some discrepancies in mitochondrial respiration results. hiPSC-derived cardiomyocytes from Barth syndrome patients or engineered to delete TAZ exhibited high basal respiration and low spare capacity (Dudek et al., 2016; Wang et al., 2014) or low basal respiration and low spare capacity (Chowdhury et al., 2023; Sniezek Carney et al., 2024). These studies, unlike ours, did not employ any maturation treatment. Isolated heart mitochondria from TAZ conditional KO mice (Zhu et al., 2021) and Barth syndrome patient mutation knock-in mice (Chowdhury et al., 2023) exhibited low basal respiration. Our TAZ KO hiPSC-derived cardiomyocytes exhibited high basal respiration and low spare capacity when cultured with maturation treatments (**Figure 1C, 2D, 5C, 7D**). While it should be carefully considered given significant phenotypic variation of TAZ knockout mice with different genetic backgrounds (Wang et al., 2023), impaired cardiac functions have been observed across different TAZ-deficient models. hiPSC-derived cardiomyocytes from Barth syndrome patients have shown contractile abnormality (Wang et al., 2014). TAZ KO mice (Wang et al., 2020; Zhu et al., 2021) and patient mutation knock-in mice (Chowdhury et al., 2023) display echocardiographic abnormalities and increased NPPA and NPPB expression. In this study, we observed TAZ KO hiPSC-derived cardiomyocytes exhibiting phenotypic variation between the two parental lines (**Figure 6, 7**). While both of our lines showed impaired contractile ability, the extent of impairment varied. High proBNP secretion was observed in one parental line but not the other. These observations highlight phenotypic variability in TAZ deficiency, which indeed mirrors Barth syndrome patients’ phenotypic spectrum (Clarke et al., 2013; Taylor et al., 2022). These discrepancies may reflect the influence of genetic modifiers associated with different parental backgrounds. Further investigation will be required to elucidate the interplay between TAZ deficiency and genetic modifiers in mitochondrial development and cardiac phenotypic variation.

The high basal respiration observed in TAZ-deficient cardiomyocytes may be attributed to increased uncoupling or proton leak. Elevated uncoupling has been implicated not only in TAZ KO mouse hearts (Wang et al., 2023) but also in other contexts of cardiac dysfunction, including diabetes (Boudina and Abel, 2006). Interestingly, dysregulated cardiolipin levels have been observed in diabetic hearts (Han et al., 2005, 2007), indicating that cardiolipin needs to be properly regulated to ensure mitochondrial integrity for optimal cardiac functionality. The detailed mechanisms underlying this phenomenon require further investigation.

In conclusion, our findings reinforce the indispensable role of cardiolipin in mitochondrial integrity. The perturbation of cristae dynamics in TAZ-deficient cardiomyocytes emphasizes the significance of TAZ-mediated cardiolipin remodeling in cardiac development. By employing the cardiomyocyte maturation system to model aspects of late fetal/neonatal cardiac transition in a dish, we identified an essential role for cardiolipin remodeling in a maturation-associated change in mitochondria that we call cardiac mitomorphosis, providing novel insights into heart development and the disease mechanisms of Barth syndrome. These discoveries may also have broader implications for understanding other cardiac dysfunctions associated with cardiolipin dysregulation (Garcia et al., 2020; Han et al., 2005, 2007; Sparagna et al., 2007).

## Limitations of this study

Our proposed model of cristae dynamics associated with cardiomyocyte maturation is only supported by static electron microscopy images. Future studies incorporating live cell imaging techniques would provide a more comprehensive understanding of these processes. Furthermore, our findings are derived from *in vitro* cardiomyocyte maturation models. Extending these investigations to track cristae dynamics during *in vivo* heart development would enhance the translational relevance of our findings.

In this study, we focused on the effects of maturation stimuli crucial for early heart development, particularly the period surrounding birth. Consequently, our findings may be most relevant to this early stage of cardiomyocyte maturation and may not fully capture later stages of development. While our maturation strategy successfully induced several hallmarks of cardiomyocyte maturation, it should be noted that the resulting cells did not achieve a fully adult-like phenotype. Unlike adult cardiomyocytes (Guo and Pu, 2020; Karbassi et al., 2020), our cardiomyocytes with maturation treatments retained a spherical morphology and lacked adult-like organized sarcomere structure, precise alignment of mitochondria, and transverse tubules. Achieving a more mature cardiomyocyte phenotype in a dish may be challenging with currently available technology, although emerging research suggests that incorporating strategies such as chronic pacing, extracellular matrix modifications, and 3D culture systems (Guo and Pu, 2020), could be beneficial.

## Acknowledgments

We thank Hilary Vernon, Ariana Anzmann, and Olivia Sniezek (Johns Hopkins University) for providing hiPSCs and valuable insights. We also thank Anastasia Kralli (Johns Hopkins University) for the generous gifts of plasmids. We acknowledge Michael Delannoy at the Johns Hopkins School of Medicine Microscopy Facility for assistance with electron microscopy. This work was supported in part by NIH grants (R01HL165729 to SMC; K99HL168075 to NS), a post-doctoral fellowship from the American Heart Association and the Barth Syndrome Foundation (Award ID: 828058 to NS), Magic that Matters Fund, The Johns Hopkins University Catalyst and Mirowski Awards (to ET), MSCRF awards (2023- MSCRFL-5984 and 2024-MSRFD-6393 to ET), and the Austrian Science Fund (FWF) projects (10.55776/P33333, 10.55776/P34574, and 10.55776/FG-15 to MAK).

## Materials and Methods

### Cell culture

#### iPSC culture

This study used two human iPSC lines: GM26105 (Coriell Institute; White male, derived from lymphocytes) and C11 (UCSFi001-A, RRID:CVCL_Y803; Asian male, derived from skin). iPSCs were maintained on Geltrex-coated plates (1:200 dilution; Gibco #A1413202) in either TeSR™-E8™ (Stem Cell Technology #05990) or mTeSR™ Plus (Stem Cell Technology #100-0276) media. iPSCs were passaged at 80-90% confluency using Accutase (Innovative Cell Technologies #AT-104). Dissociated cells were collected with DMEM supplemented with 10% FBS (Sigma #F2442), resuspended in TeSR™-E8™ or mTeSR™ Plus containing RevitaCell™ (Gibco #A2644501) or CloneR™2 (Stem Cell Technology #100-0691), and plated at a density of 15,000-20,000 cells/cm^2^.

#### iPSC-cardiomyocyte culture

Cardiomyocyte differentiation from human iPSCs was performed based on the previously described canonical Wnt stimulation and inhibition protocol (Maas et al., 2021). iPSCs were plated on Geltrex-coated 12-well plates (1:200 dilution) and grown to 80-90% confluency for differentiation. On day 0, iPSCs were treated with 4 μM CHIR99021 (Tocris #4423) in RPMI 1640 supplemented with B27 without insulin (Gibco #A1895602) for 24 hours. On day 1, the medium was replaced with 2 μM CHIR in RPMI 1640 supplemented with B27 minus insulin for 48 hours. On day 3, the medium was replaced with 5 μM IWR-1 (Sigma-Aldrich #I0161) in RPMI 1640 supplemented with B27 minus insulin for 48 hours. On day 5, the medium was replaced with RPMI 1640 supplemented with B27 minus vitamin A (Gibco #12587001). The medium (RPMI 1640 supplemented with B27 minus vitamin A) was replaced every two days until days 9-11 when spontaneous contractions began to be observed. The medium was then replaced with glucose-free RPMI 1640 supplemented with 5 mM lactate (Sigma #71718) for 2 days. By this point, the majority of cells were spontaneously contracting cardiomyocytes. The medium was then replaced with RPMI 1640 supplemented with B27 minus vitamin A for 2-4 days to allow for recovery. On days 15-18, cardiomyocytes were dissociated using TrypLE Select 10X (Gibco #A12177) and either replated or cryopreserved in STEMdiff™ Cardiomyocyte Freezing Medium (Stem Cell Technology #05030). Replated or thawed iPSC-derived cardiomyocytes were settled in RPMI 1640 supplemented with B27 minus vitamin A and 10% FBS (Sigma) on 0.1% gelatin-coated plates for 1-2 days, then expanded in RPMI 1640 supplemented with B27 minus vitamin A containing 2 μM CHIR99021, as previously described (Maas et al., 2021), until approximately 30 days post-differentiation initiation. The expanded cardiomyocytes were cultured in RPMI 1640 supplemented with B27 minus vitamin A with or without the described treatments.

#### Compounds for maturation treatments

Palmitic acid-BSA conjugate was made as follows: Palmitic acid (Sigma #P0500) was dissolved in 90 mM NaOH to a final concentration of 100 mM and solubilized by incubation at 90°C for 10 minutes. Fatty acid-free BSA (Sigma #A7030) was dissolved in 0.1 M Tris to a final concentration of 3.84 mM and warmed to 37°C. The palmitic acid solution was then diluted to 20 mM with the BSA solution and vortexed until completely dissolved. The 20 mM palmitic acid-BSA conjugate was diluted in RPMI 1640 to 1 mM to prepare stock solution. The diluted palmitic acid-BSA conjugate was added to RPMI 1640 supplemented with B27 minus vitamin A to a final concentration of 20 μM. L-carnitine (Sigma #C0283), dexamethasone (Sigma-Aldrich #D4902), and triiodothyronine (Sigma-Aldrich #T6397) were purchased and dissolved in RPMI 1640 to prepare stock solutions. These were added to the culture media at final concentrations of 2 mM (L-carnitine), 1 μM (dexamethasone), and 32 ng/ml (triiodothyronine), respectively.

#### Endothelin-1 treatment

Endothelin-1 (ET-1; Sigma #E7764) was dissolved in RPMI 1640 to prepare a 10 μM stock solution. This stock solution was added to RPMI 1640 supplemented with B27 minus vitamin A, Pal/car, and Dex/T3 for a final ET-1 concentration of 10 nM. iPSC-derived cardiomyocytes were cultured in this medium for 2 weeks.

Mycoplasma contamination was routinely monitored using Universal Mycoplasma Detection Kit (ATCC #30-1012K). Sterility was maintained throughout all experiments using standard aseptic techniques.

### Knockout iPSC lines

TAZ KO in GM26105 iPSC line was previously generated using CRISPR/Cas9 genome editing, as described in (Sniezek Carney et al., 2024). This study employed the same strategy to generate TAZ KO C11 iPSC lines. Briefly, two guide RNAs were designed to remove 50 bp at the conserved acyltransferase domain of the human *TAFAZZIN* gene, and cloned into pSpCas9(BB)-2APuro (PX459) V2.0, a gift from Feng Zhang (Addgene plasmid #62988). Two guide RNA vectors were transfected into C11 iPSCs using Lipofectamine Stem (Invitrogen #STEM00001). Transfected cells were selected with puromycin (Gibco #A11138) at 0.25 μg/ml for 48 hours. Single-cell clones were then isolated by limiting dilution on a 96- well plate. Individual clones were expanded and screened for TAZ knockout by immunoblot analysis. For genotyping, genomic DNA was extracted from KO clones using Gentra Puregene Cell Kit (Qiagen). The region flanking the gDNA target sites was amplified by PCR, using the following primers (forward, 5’- GGAATTCGGATGCCTCTGCACGTGAAG-3’; reverse, 5’-CGGGATCCAAACCGGAGTGCCAGGGCTC-3’), and cloned into the pBSK(-) vector. Transformants were analyzed by Sanger sequencing.

### Lentivirus transduction

WT TAZ or H69Q mutant, as we previously reported (Lu et al., 2016), were subcloned into the pFUGW backbone plasmid (EF-1α promoter) by NEBuilder HiFi assembly (NEB #E2621S). The primers used for cloning are: Forward, 5’-TTTGCCGCCAGAACACAGGACCGGTAAACTTAAGCTTATGCCTC-3’; reverse, 5’-GAGAGAAGTTTGTTGCGCCGGATCCTTTAAACGGGCCCTCTAG-3’. 293T cells were plated on 10 cm dishes and grown to 50-70% confluency. The respective pFUGW TAZ plasmid was co-transfected with the packaging plasmids psPAX2 and pMD2.G using 1 mg/ml PEI solution (Polysciences #24765). 24 and 48 hours post-transfection, the culture media were collected, filtered, and supplemented with polybrene (4 μg/ml final concentration; Sigma #TR-1003). For transduction, TAZ knockout iPSCs (GM26105 iPSC line) were incubated with the respective lentivirus-containing media. After 24 hours, the media was replaced with TeSR™-E8™. Cells were then selected with 0.5 μg/ml puromycin for 48 hours. Single-cell clones were isolated by limiting dilution on a 96-well plate. Individual clones were expanded and screened for TAZ expression by immunoblot analysis. iPSC clones that expressed the transduced TAZ at a similar level to endogenous TAZ were selected for the subsequent experiments.

### Cardiomyocyte purity

iPSC-derived cardiomyocytes were dissociated using Accumax (Innovative Cell Technologies #AM105) and collected in PBS. For fixation, cells were pelleted by centrifugation at 1000 g for 3 minutes, then resuspended in 1% paraformaldehyde (Thermo Scientific Chemicals #J61899-AK) diluted in PBS and incubated for 20 minutes. Following another centrifugation at 1000 g for 3 minutes, cells were resuspended in ice-cold 90% methanol in PBS and incubated for 15 minutes on ice for permeabilization.

Cells were then washed twice with 0.5% BSA (Sigma #A9647) in PBS, followed by incubation with a primary antibody against cardiac Troponin T (cTnT, 2 µg/mL; mouse monoclonal (RV-C2), Developmental Studies Hybridoma Bank) in 0.5% BSA in PBS with 0.1% Triton X-100 for 2 hours. After a single wash with 0.5% BSA in PBS containing 0.1% Triton X-100, cells were incubated with a secondary antibody (1:500; DyLight 650 goat anti-mouse IgG (H+L), Invitrogen #84545) in 0.5% BSA in PBS with 0.1% Triton X-100 containing 1 µg/mL Hoechst 33342 (Thermo Scientific #62249) for 30 minutes. Cells were then washed twice with 0.5% BSA in PBS containing 0.1% Triton X-100, resuspended, and diluted in 0.5% BSA. Cell dilutions were plated on a black-wall 96-well plate and scanned using a Celigo Cytometer (Nexcelom). Cardiomyocyte purity was determined as the percentage of cTnT-positive cell population (identified by cTnT staining) relative to the total cell population (identified by Hoechst staining).

### Cell viability

iPSC-derived cardiomyocytes were plated at a density of 40,000 cells/well in a black-walled 96-well plate and cultured with the respective treatments. Cell viability was assessed by staining with ViaStain AOPI Staining Solution (Nexcelom #CS2-0106) supplemented with 1 µg/mL Hoechst 33342 (Thermo Scientific #62249). Cells were incubated with the staining solution in a 37°C CO2 incubator for 15 minutes. Following a single wash with PBS containing 2% FBS, cells were scanned using a Celigo Cytometer (Nexcelom). The percentage of PI-positive cells within the total cell population (identified by Hoechst staining) was quantified as dead cells.

### SDS-PAGE and immunoblot

Cell pellets were lysed in RIPA buffer (1% (v/v) Triton X-100, 20 mM HEPES-KOH, pH 7.4, 50 mM NaCl, 1 mM EDTA, 2.5 mM MgCl2,0.1% (w/v) SDS) spiked with protease inhibitors (1 mM PMSF, 2 μM pepstatin A, 10 μM leupeptin). Protein concentrations were determined using Pierce BCA Protein Assay Reagents (Thermo Scientific™ #23228, #23228). Lysates were mixed with reducing sample buffer (20% (w/v) glycerol, 4% (w/v) SDS, 0.2M DTT, Tris-Cl), incubated at 95°C for 5 minutes, and loaded on Tris-glycine gels for SDS-PAGE. Proteins were transferred to nitrocellulose membranes (0.45 μm, Bio-Rad). Membranes were blocked in 5% skim milk in PBST (PBS with 0.05% Tween 20) and incubated with primary antibodies diluted in PBST, followed by incubation with HRP-conjugated secondary antibodies diluted in PBST. Clarity Western ECL Substrate (Bio-Rad #170-5061) was applied to the membranes, and chemiluminescence was detected using a ChemiDoc MP Imaging System (Bio-Rad). Antibodies used in this study are as follows: TAZ (mouse monoclonal (2C2C9) (Lu et al., 2016)), CPT1B (rabbit polyclonal, Proteintech #22170-1-AP), CPT2 (rabbit monoclonal (EPR13626), Abcam #ab181114), MPC1 (rabbit monoclonal (D2L9I), Cell Signaling Technology #14462), MPC2 (rabbit monoclonal (D417G), Cell Signaling Technology #46141), NDUFB6 (mouse monoclonal (21C11BC11), Abcam #ab110244), COX4 (rabbit polyclonal, Abcam #ab16056), UQCRC2 (mouse monoclonal (13G12), Abcam #ab14745), MT- CO1 (mouse monoclonal (1D6E1A8), Thermo Fisher Scientific #459600), SDHA (mouse monoclonal (2E3GC12FB2AE2), Abcam #ab14715), F1β (ATP5B) (rabbit polyclonal, Proteintech #17247-1-AP), TOM20 (rabbit polyclonal, Proteintech #11802-1-AP), cTnI (mouse monoclonal (TI-1), Developmental Studies Hybridoma Bank), ssTnI (mouse monoclonal (TI-4), Developmental Studies Hybridoma Bank), ANT1 (mouse monoclonal (1F3F11), (Lu et al., 2017)), ANT2 (rabbit polyclonal, (Senoo et al., 2024)), HRP-conjugated secondary, goat anti-rabbit IgG (H+L) (Thermo Fisher Scientific #31460), HRP-conjugated secondary, goat anti-mouse IgG (H+L) (Thermo Fisher Scientific #62-6520).

### Cardiolipin measurements by LC-MS/MS

Lipid extraction, mass spectrometric cardiolipin analysis and relative quantification were carried out as detailed in (Wohlfarter et al., 2022). Briefly, with cardiolipins being exclusively localized to mitochondria, no subcellular fractionation was necessary. Lipids were extracted using the Folch method (Folch et al., 1957) and subsequently separated by reversed-phase high-performance liquid chromatography (RP-HPLC). Lipid detection was conducted using time-of-flight mass spectrometry (timsTOF Pro, Bruker Daltonics, Bremen, Germany). Raw data were processed and analyzed via a targeted feature integration workflow implemented in MZmine 2.53 (Giombini et al., 2010), followed by manual curation of all relevant signals.

### Seahorse analysis

After the respective treatment, iPSC-derived cardiomyocytes were dissociated using TrypLE Select (diluted to 5X) and seeded onto XF96e cell culture microplates coated with 0.1% gelatin in RPMI 1640 supplemented with B27 minus vitamin A, 10% FBS, and RevitaCell. Plating cell densities were as follows: GM26105 parental cells, 8-day Pal/car treatment: 100,000 cells/well; GM26105 parental cells, 2-week Pal/car + Dex/T3 treatment: 80,000 cells/well; C11 parental cells, 2-week Pal/car + Dex/T3 treatment: 40,000 cells/well. The day after plating, the medium was replaced with the respective treatment medium, and the cells were incubated for 2 days. On the day of measurement, cells were washed twice and then incubated for 1 hour in a 37°C non-CO2 incubator in Seahorse XF DMEM Basal Medium (Agilent #103334-100) supplemented with 10 mM glucose (Sigma #G8644), 2 mM L-glutamine (Gibco #25030-081), 1 mM sodium pyruvate (Gibco #11360-070), 0.5 mM L-carnitine (Sigma #C0283), and 160 μM BSA-conjugated palmitic acid (Cayman #29558). MitoStress Test (Agilent #103015-100) was performed on Seahorse XFe96 Analyzer (Agilent) by injecting oligomycin (2.5 μM), FCCP (1 μM), rotenone and antimycin A (0.5 μM) sequentially. To assess the reliance on pyruvate or fatty acid oxidation, after recording baseline oxygen consumption rate, UK-5099 (30 μM; Tocris #4186) or etomoxir (200 μM; Tocris #4539) was injected. The acute response to each inhibitor was determined as the change in oxygen consumption rate following inhibitor injection.

### Transmission electron microscopy

iPSC-derived cardiomyocytes were cultured on 0.1% gelatin-coated 35 mm dishes in the respective treatment media. Cells were fixed with 2% glutaraldehyde (Electron Microscopy Sciences #16220) in 0.1 M cacodylate/3 mM CaCl2, washed three times with 0.1 M cacodylate/3 mM CaCl2, and postfixed with 1% OsO4 (Electron Microscopy Sciences #19190) reduced with potassium hexacyanoferrate (Sigma-Aldrich #P3289) in 0.1 M cacodylate. Fixed cells were stained with 2% uranyl acetate (Ted Pella, Inc. #19481), dehydrated in a graded series of ethanol, and embedded in Epon (EMBED 812 RESIN, DDSA, NMA; Electron Microscopy Sciences #14900, #13710, #19000) with 1.5% DMP-30 catalyst (Electron Microscopy Sciences #13600). Ultrathin sections were examined using a Hitachi 7600 transmission electron microscope.

Sample preparation for transmission electron microscopy was performed for at least 3 biological replicates. Fixation and imaging were done by separate individuals. Quantitative analyses were performed in a blinded manner, with sample groups randomized to prevent bias during image analysis. To determine cristae detachment, the number of cristae detached from the IBM per mitochondrion was counted manually. Cristae were considered detached if there was a clear separation from the IBM. The number of detached cristae was normalized to mitochondrial perimeter measured using ImageJ. To assess morphological variation, mitochondria were categorized into 4 metrics based on their appearance: defined (>50% cristae with clear and aligned membranes), intermediate (25-50% cristae with clear and aligned membranes), irregular (<25% cristae with clear and aligned membranes), and rupture. The proportion of mitochondria in each category was calculated for each biological replicate. To determine cristae density, the length of cristae membranes was traced manually using ImageJ. The sum of cristae membrane length per mitochondrion was normalized to mitochondrial perimeter.

### Blue native-PAGE

iPSC-derived cardiomyocyte pellets were pre-treated with 0.4% digitonin in PBS for 10 minutes on ice according to a previously published protocol (Timón-Gómez et al., 2020). The pellets were then centrifuged at 1,000 g for 5 minutes at 4°C and washed with PBS. The washed pellets were homogenized in 100 mM Bis-Tris, pH 7.0/1 M 6-aminocaproic acid supplemented with protease inhibitors (1 mM PMSF, 2 μM pepstatin A, 10 μM leupeptin). Protein concentration was measured with Pierce BCA Protein Assay Reagents (Thermo Scientific™). For Complexes I, III and IV analysis, homogenates (200 µg) were mixed with Triton X-100 (protein:Triton X-100 = 1:4) or n-dodecyl-β-D-maltoside (DDM) (protein:DDM = 1:4). For respiratory supercomplexes and Complex V analyses, homogenates (200 µg) were mixed with digitonin (protein:digitonin = 1:1, 1:2, or 1:4). Samples were solubilized on ice for 30 minutes. Following solubilization, samples were centrifuged at 21,000 g for 30 minutes at 4°C. The supernatants were collected and mixed with 10X blue native-PAGE sample buffer (5% (w/v) Coomassie Brilliant Blue G- 250 (Serva), 0.5 M 6-aminocaproic acid, and 10 mM Bis-Tris, pH 7.0). The extracts were loaded on house-made blue native-PAGE gels. Electrophoresis was performed at 100-120V in a cold room. Blue cathode buffer (50 mM Tricine, 15 mM Bis-Tris, pH 7.0) containing Coomassie Brilliant Blue G-250 (Serva) was used initially, then replaced with clear cathode buffer, and running was performed until the dye front reached the bottom. Proteins were transferred to PVDF membranes (0.45 μm, Millipore), and immunoblotting was performed as described above.

### mRNA expression

Total RNA was extracted from iPSC-derived cardiomyocytes using the PureLink RNA Mini Kit (Invitrogen #12183018A) according to the manufacturer’s instructions. cDNA was synthesized using SuperScript VILO™ (Invitrogen #11755-050). qPCR was performed using Fast SYBR Green Master Mix (Applied Biosystems #4385612) on a Rotor-Gene Q instrument (Qiagen). The reaction was performed as follows: initial denaturation at 95°C for 20 seconds, followed by 40 cycles of denaturation at 95°C for 1 second and annealing/extension at 60°C for 20 seconds. Subsequently, melt curve analysis was performed. Relative gene expression was analyzed using the ΔΔCT method, with EEF1A1 or YWHAZ as reference genes. Primer sequences are listed in Table S1.

### Mitochondrial DNA copy number

Genomic DNA was extracted from iPSC-derived cardiomyocytes using the Gentra Puregene Cell Kit (Qiagen). qPCR was performed as described above (see ’mRNA analysis’ section) using primers for MT- ND1 (mitochondrial DNA-encoded) and B2M (nuclear DNA-encoded) (listed in Table S1). MT-ND1 expression was normalized to B2M.

### proBNP secretion

iPSC-derived cardiomyocytes were plated onto a 24-well plate at 400,000 cells/well and cultured for 2 weeks in the respective treatment medium. Medium change (1 ml/well) was performed every 3-4 days. The last medium change was done 4 days before supernatant collection. The supernatants were collected and stored at -80°C until use. proBNP levels in the supernatant were measured using the Human proBNP ELISA kit (Invitrogen #EHPRONPPB) according to the manufacturer’s instructions.

### Contraction assessment

iPSC-derived cardiomyocytes were plated onto a 24-well plate at 400,000 cells/well and cultured for 2 weeks with medium changes every 3-4 days. On the day of the assay, cells were fed with fresh RPMI 1640 supplemented with B27 minus vitamin A, Pal/car, and Dex/T3 and incubated for 1 hour at 37°C. A fresh isoproterenol stock solution (100 mM) was prepared by dissolving isoproterenol (Sigma-Aldrich #I6379) in 1 M HCl. This stock solution was then serially diluted in RPMI 1640 supplemented with B27 minus vitamin A, Pal/car, and Dex/T3 for the desired final concentrations. Cardiomyocyte contractions were video-recorded (40 fps) using a BZ-X710 microscope (Keyence). Baseline contractions were recorded for 30 seconds. Then, isoproterenol was added sequentially to achieve final concentrations of 10 nM, 100 nM, and 1 μM. After each addition of isoproterenol, the plate was gently swirled and contractions were recorded for 30 seconds. Video recordings were analyzed using the ImageJ plugin MYOCYTER to obtain parameters of contraction frequency, amplitude, and systolic and diastolic peak times, as previously described (Grune et al., 2019).

