## Supplemental Figures S1-S15 and Table S1 for "Disturbed mitochondrial maturation in cardiolipin remodeling-deficient cardiomyocytes"

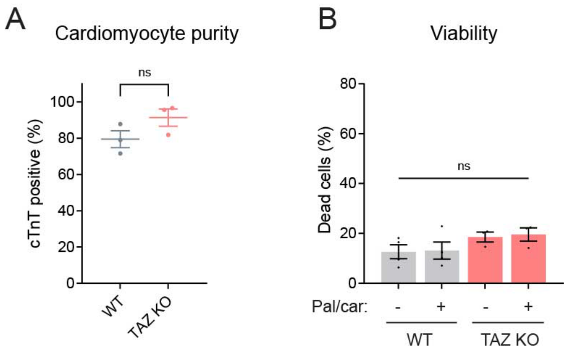


**Figure S1: Purity and viability of WT and TAZ KO iPSC-derived cardiomyocytes.**

**(A)** Cardiomyocyte purity, presented as cardiac troponin T (cTnT) positive population. WT and TAZ KO cardiomyocytes at baseline were analyzed (mean ± SEM, n = 3 biological replicates). **(B)** Viability of WT and TAZ KO cardiomyocytes after 8 days of treatment with Pal/car (mean ± SEM, n = 3-4 biological replicates).

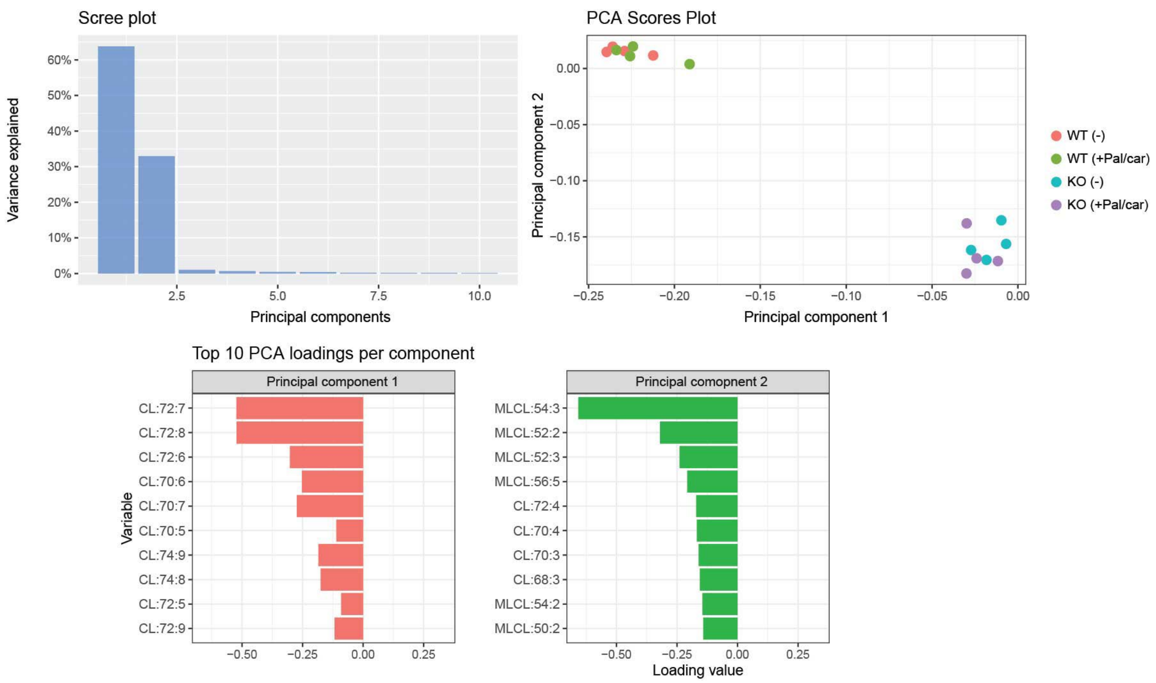


**Figure S2: Principal component analysis of cardiolipin molecular species composition in cardiomyocytes cultured under Pal/car for 8 days.**

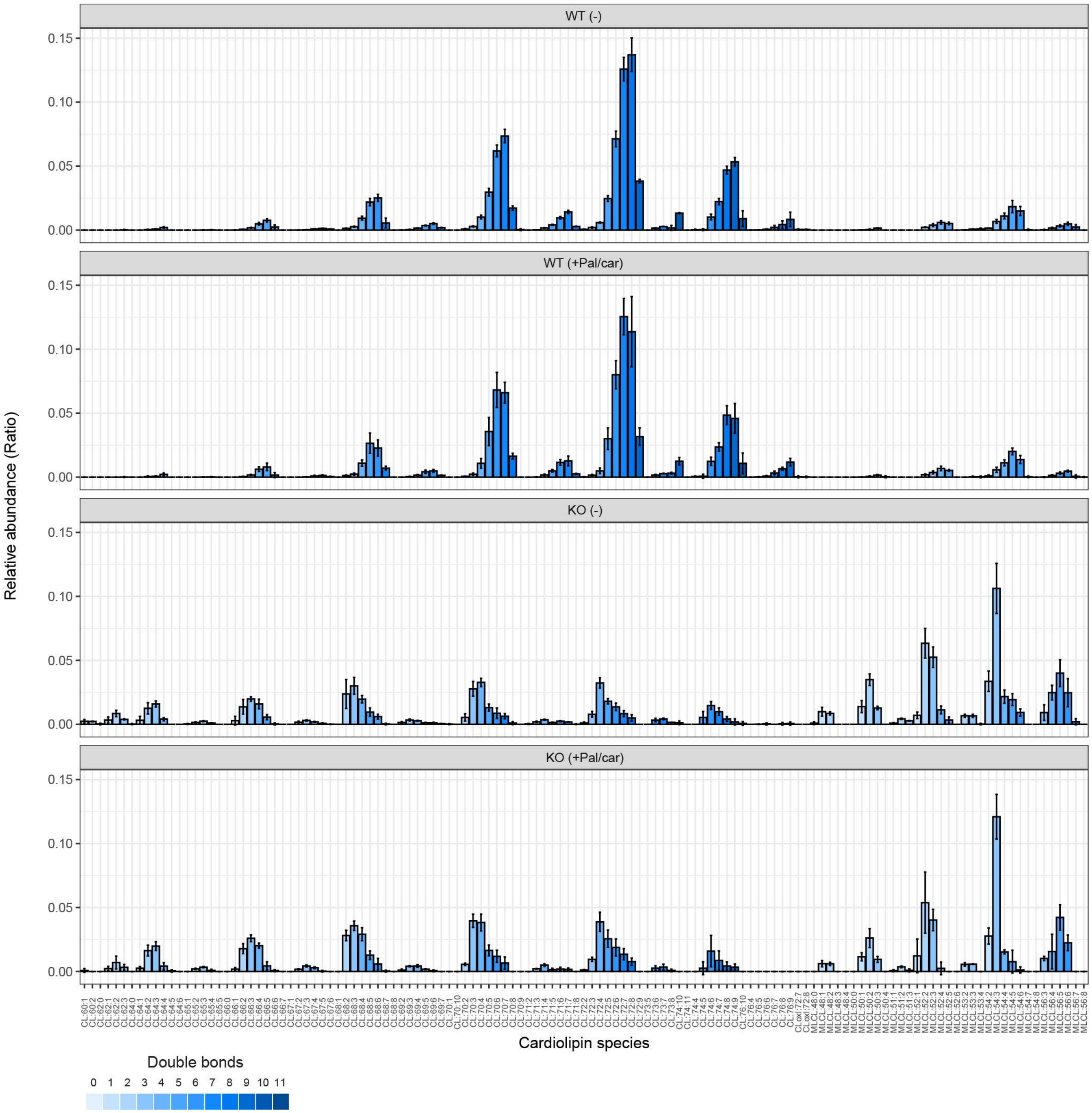


**Figure S3: Cardiolipin and monolyso-cardiolipin molecular species in cardiomyocytes cultured under Pal/car for 8 days.** Data are shown as mean ± SD (n = 4 biological replicates).

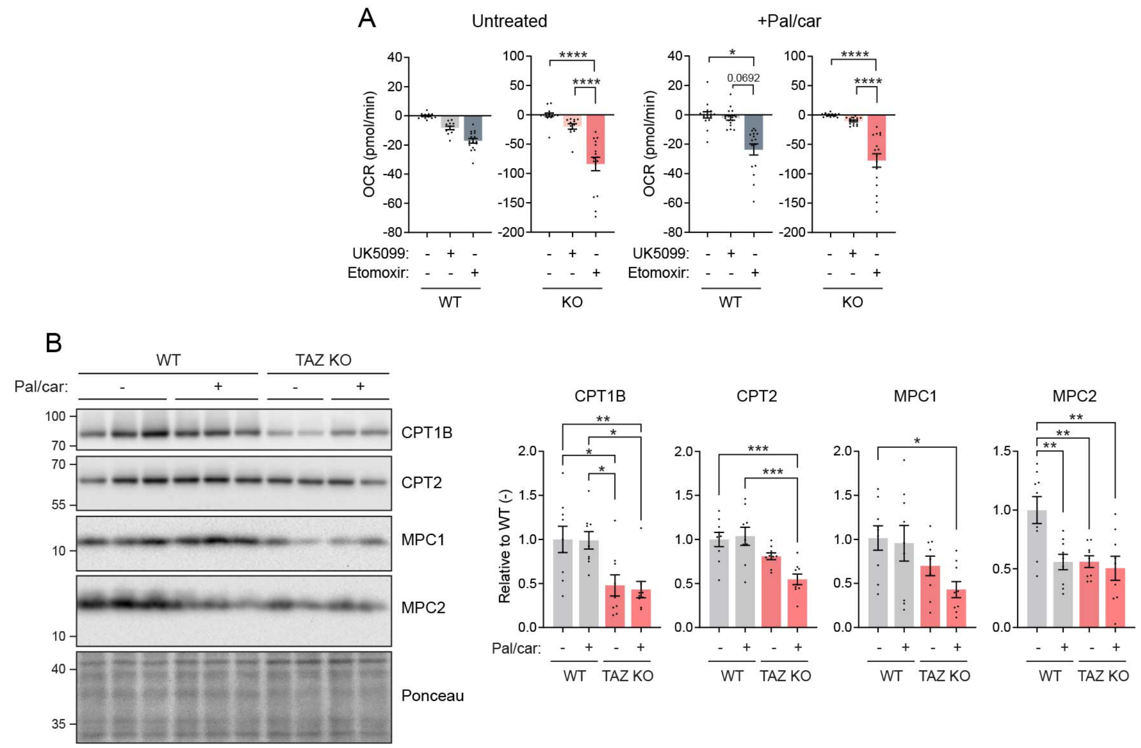


**Figure S4: Respiratory substrate reliance of cardiomyocytes cultured under Pal/car for 8 days.**

**(A)** Related to Figure 1C, OCR acute responses to the injections of UK-5099 (30 µM) or etomoxir (200 µM) were measured by Seahorse XF96e FluxAnalyzer. **(B)** Immunoblot analysis of CPT and MPC expression (mean ± SEM, n = 9 biological replicates). Statistical significance: one-way ANOVA with Tukey's test; *p<0.05, **p<0.01, ***p < 0.001, ****p < 0.0001.

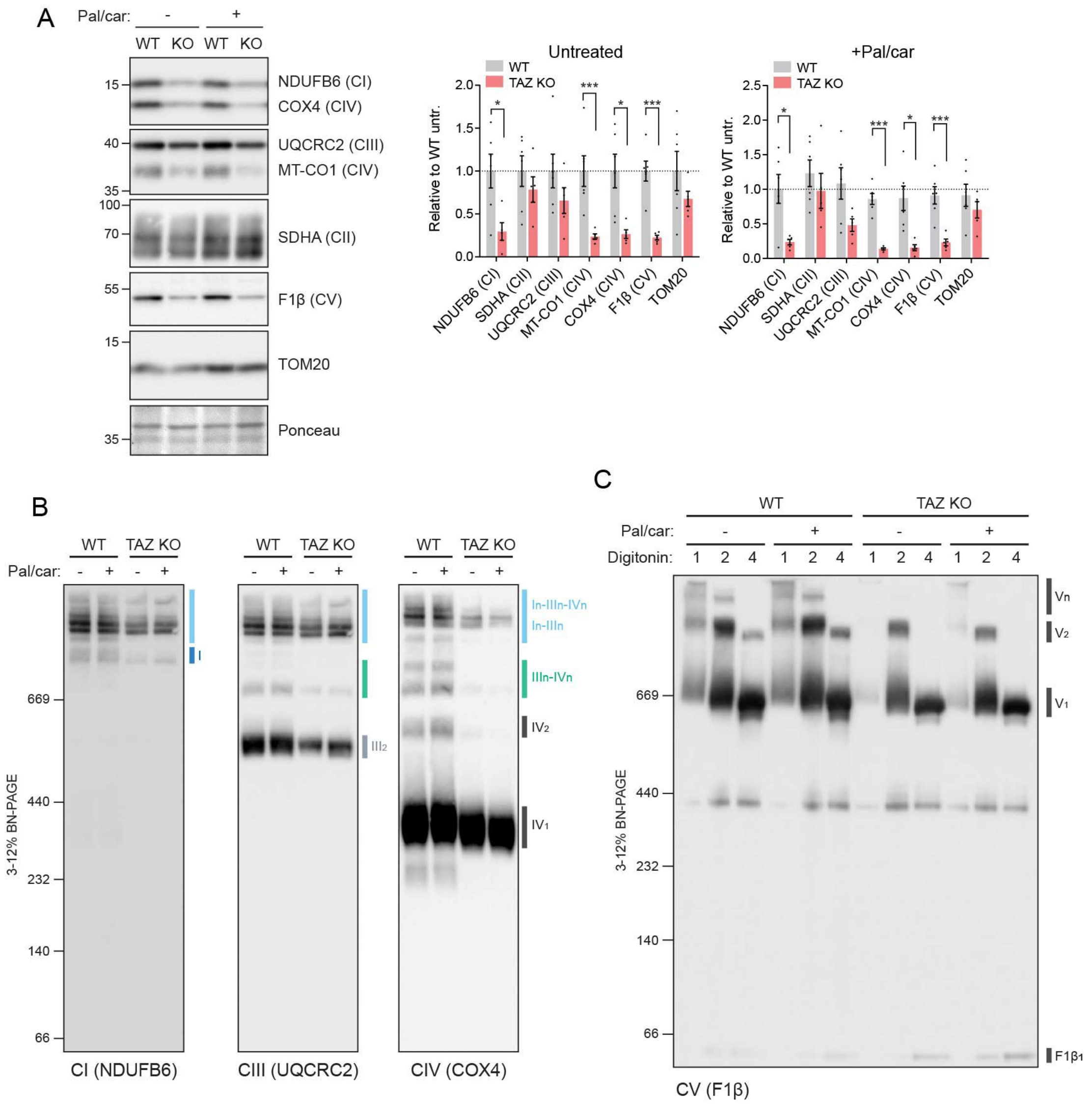


**Figure S5: OXPHOS machinery in cardiomyocytes cultured under Pal/car for 8 days.**

WT and TAZ KO cardiomyocytes were cultured with Pal/car for 8 days. **(A)** Immunoblot analysis of OXPHOS subunits in cell extracts (Complexes I, II, III, IV, and V) (mean ± SEM, n = 5-6 biological replicates). Statistical significance: one-way ANOVA with Tukey's test; *p<0.05, **p<0.01, ***p < 0.001, ****p < 0.0001. **(B)** Respiratory supercomplex assembly. Digitonin-solubilized cell extracts (protein:digitonin = 1:4) were analyzed by blue native-PAGE and immunoblot. Representative images from n = 3 biological replicates are shown. **(C)** Complex V assembly. Digitonin-solubilized cell extracts (protein:digitonin = 1:1, 1:2, or 1:4) were analyzed by blue native-PAGE and immunoblot. Numerical subscripts specify the number of individual complex units comprising the indicated assembly; 'n' denotes a variable number. A representative image from n = 3 biological replicates is shown.

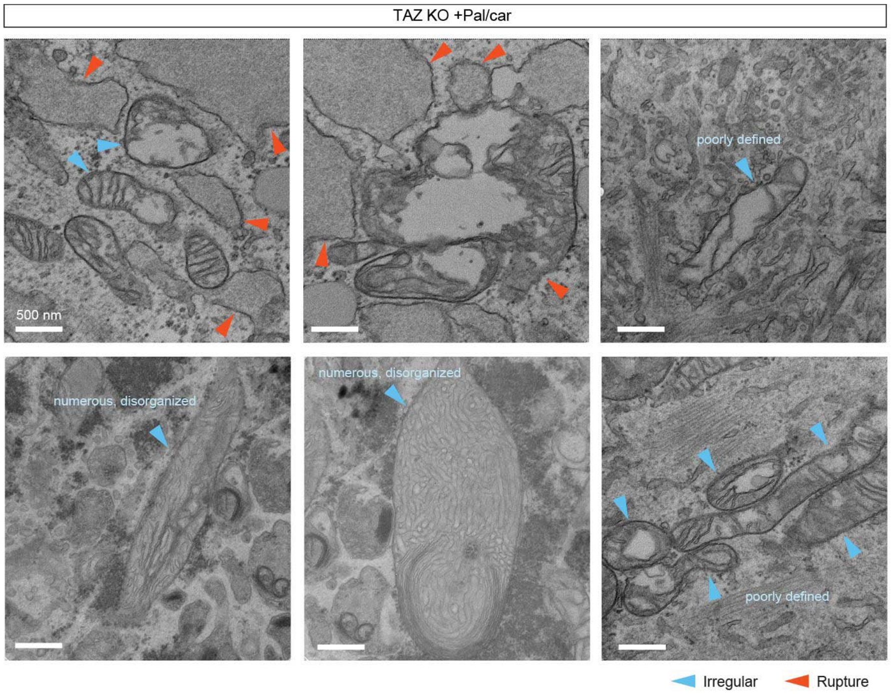


**Figure S6: Electron microscopy images of cristae morphology in TAZ KO cells with 8-day Pal/car treatment.**
Related to Figure 1D, additional images of cristae morphology in TAZ KO cells with 8-day Pal/car treatment are shown.

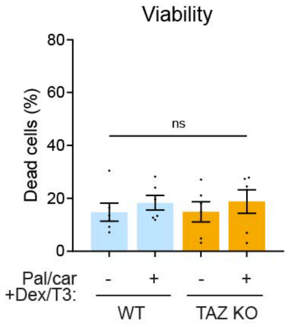


**Figure S7: Viability of WT and TAZ KO iPSC-derived cardiomyocytes cultured with hormone treatment.**

WT and TAZ KO cardiomyocytes were cultured for 2 weeks with Pal/car + Dex/T3 treatment (mean ± SEM, n = 6 biological replicates).

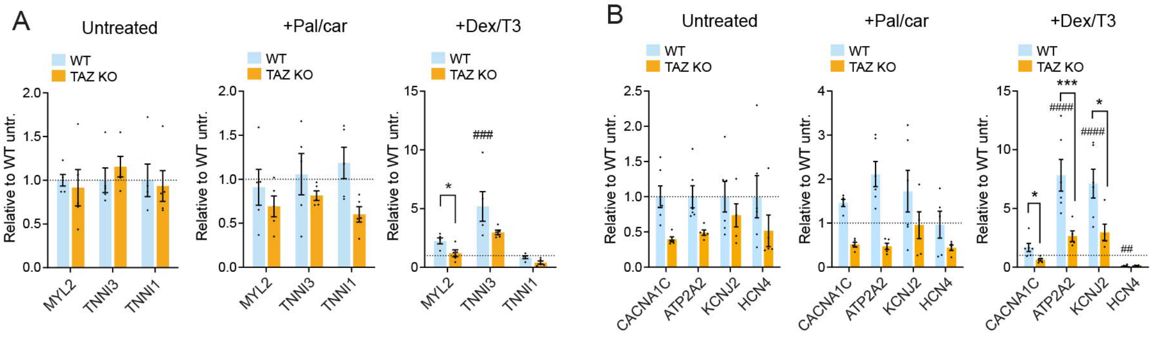


**Figure S8: Maturation marker expression in cardiomyocytes.**

**(A, B)** Related to Figure 2A and C, WT and TAZ KO cardiomyocytes were cultured with the indicated treatments for 2 weeks. Sarcomere (A) and electrophysiological (B) gene expression were analyzed by qPCR normalized to *EEF1A1* (mean ± SEM, n = 5 biological replicates). Dashed lines: untreated WT. Statistical significance: one-way ANOVA with Tukey's test; *p<0.05, **p<0.01, ***p < 0.001, ****p < 0.0001. * WT vs. TAZ KO within treatment; # vs. untreated within genotype.

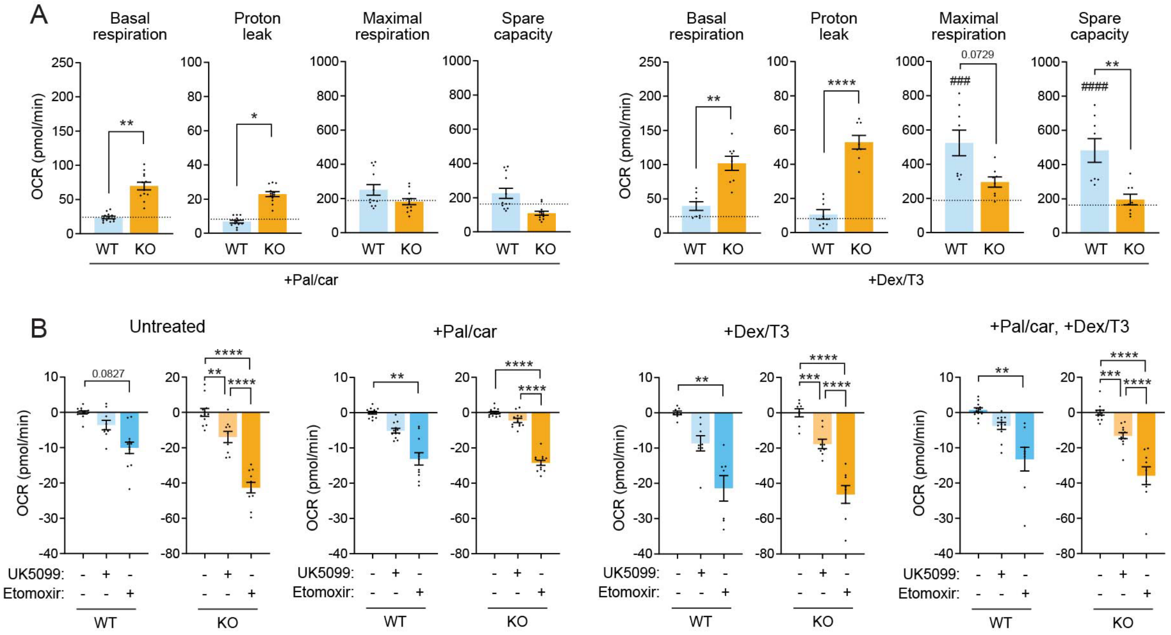


**Figure S9: Mitochondrial respiration in cardiomyocytes with 2-week treatments.**

Related to Figure 2D, WT and TAZ KO cardiomyocytes were cultured for 2 weeks with the indicated treatments. OCR was measured by Seahorse XF96e FluxAnalyzer with glucose, glutamine, pyruvate, and palmitic acid as substrates (80K/well; mean ± SEM, n = 9-12 with 3 biological replicates and 3-4 technical replicates). **(A)** Parameters obtained from the Mito Stress Test. Dashed lines: untreated WT (shown in Figure 2). **(B)** Acute responses to the injections of UK-5099 (30 μM) or etomoxir (200 μM). Statistical significance: one-way ANOVA with Tukey's test; *p<0.05, **p<0.01, ***p < 0.001, ****p < 0.0001. * WT vs. TAZ KO within treatment; # vs. untreated within genotype.

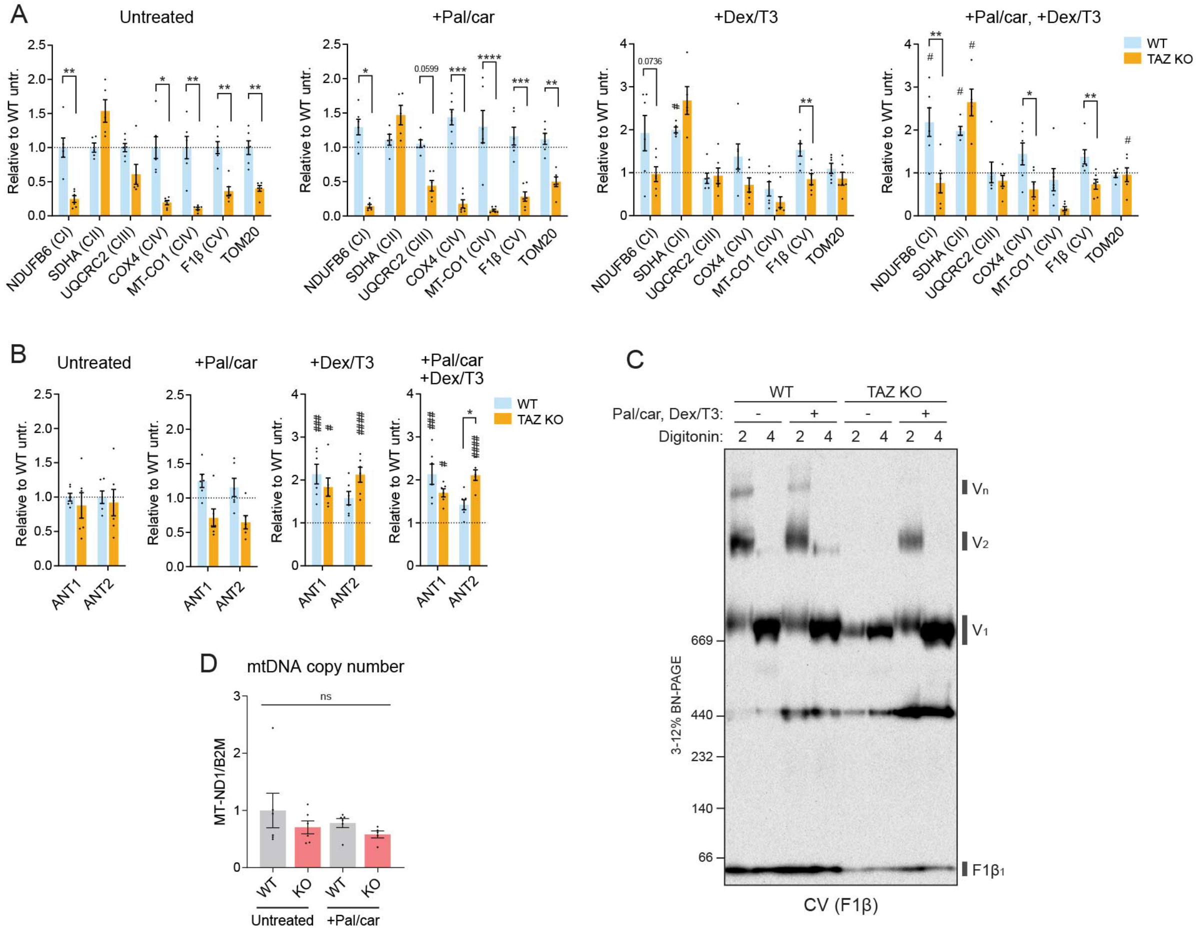

**Figure S10: Additional analysis for the inner mitochondrial membrane protein machinery in mature cardiomyocytes.**

**(A, B)** Related to Figure 3A, D, quantification of immunoblot analysis of OXPHOS protein levels in WT and TAZ KO iPSC-derived cardiomyocytes cultured with the indicated treatments for 2 weeks are shown. Dashed lines: untreated WT. **(C)** Complex V assembly in WT and TAZ KO cardiomyocytes cultured with the indicated treatments for 2 weeks. Digitonin-solubilized cell extracts (protein:digitonin = 1:2 or 1:4) were analyzed by blue native-PAGE and immunoblot. A representative image from n = 3 biological replicates is shown. **(D)** Mitochondrial DNA copy number analyzed by qPCR in cardiomyocytes cultured with or without Pal/car for 8 days (mean ± SEM, n = 5-6 biological replicates). Statistical significance: one-way ANOVA with Tukey's test; *p<0.05, **p<0.01, ***p < 0.001, ****p < 0.0001. * WT vs. TAZ KO within treatment; # vs. untreated within genotype.

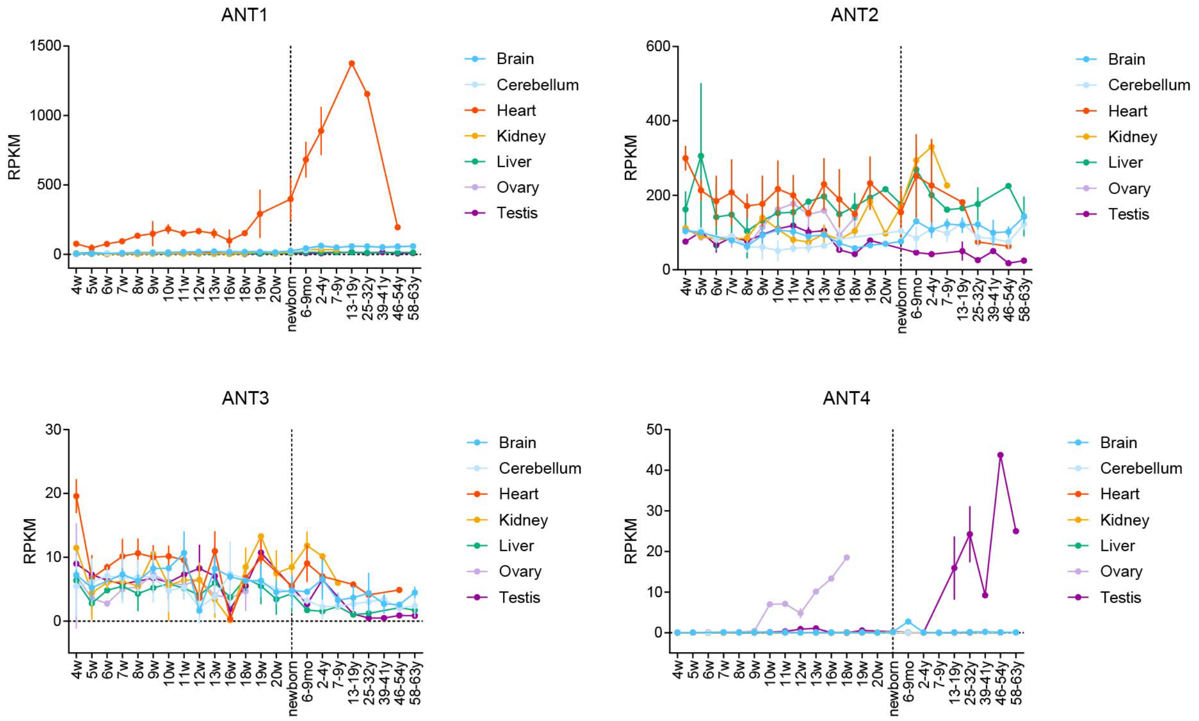


**Figure S11: ANT isoform expression across organs and developmental stages.**

RPKM of human ANT isoforms obtained from the Evo-devo mammalian organ expression atlas (Cardoso-Moreira et al., 2019).

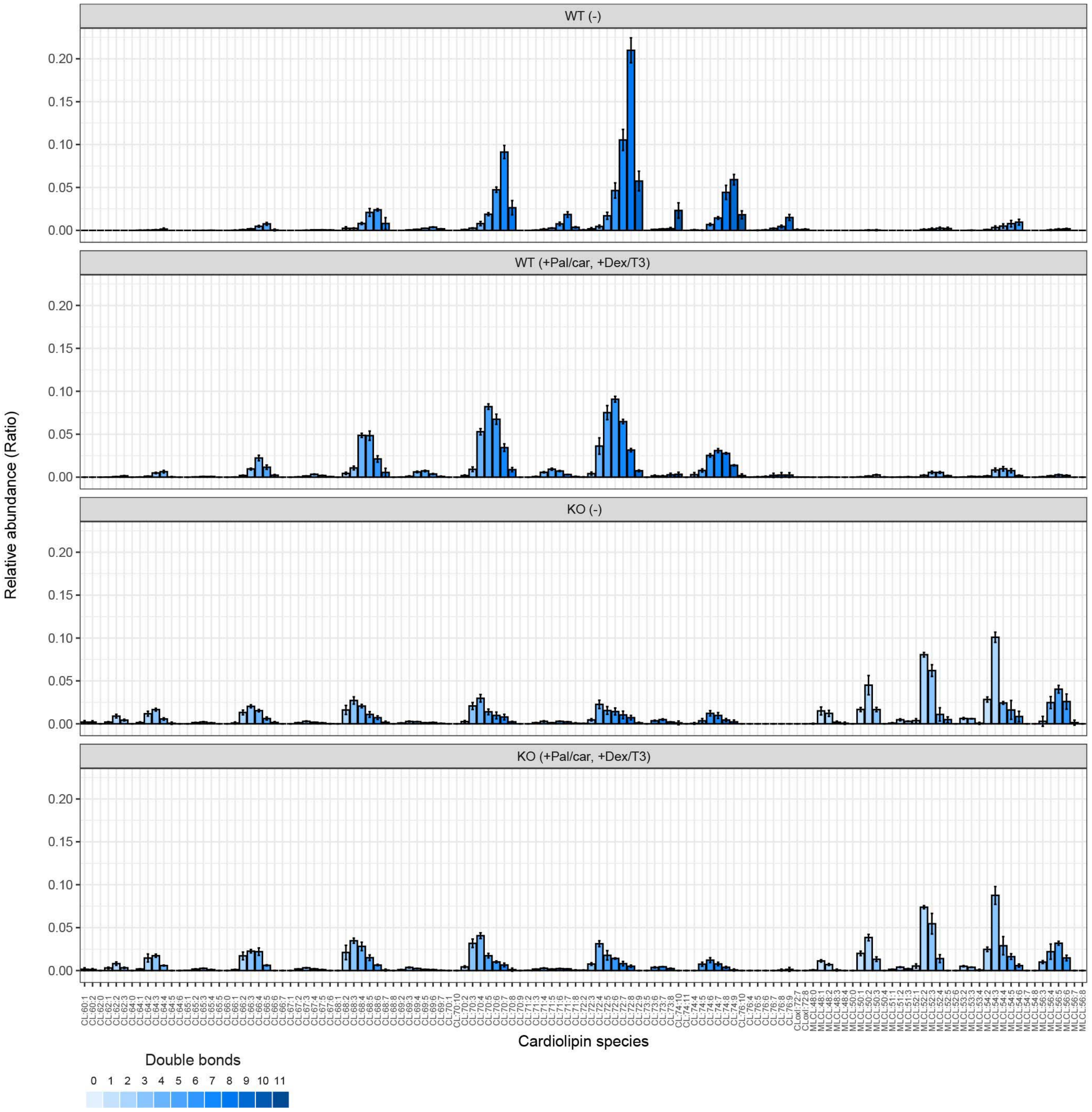


**Figure S12: Cardiolipin and monolyso-cardiolipin molecular species in cardiomyocytes cultured under Pal/car + Dex/T3 for 2 weeks.** Data are shown as mean ± SD (n = 3-4 biological replicates).

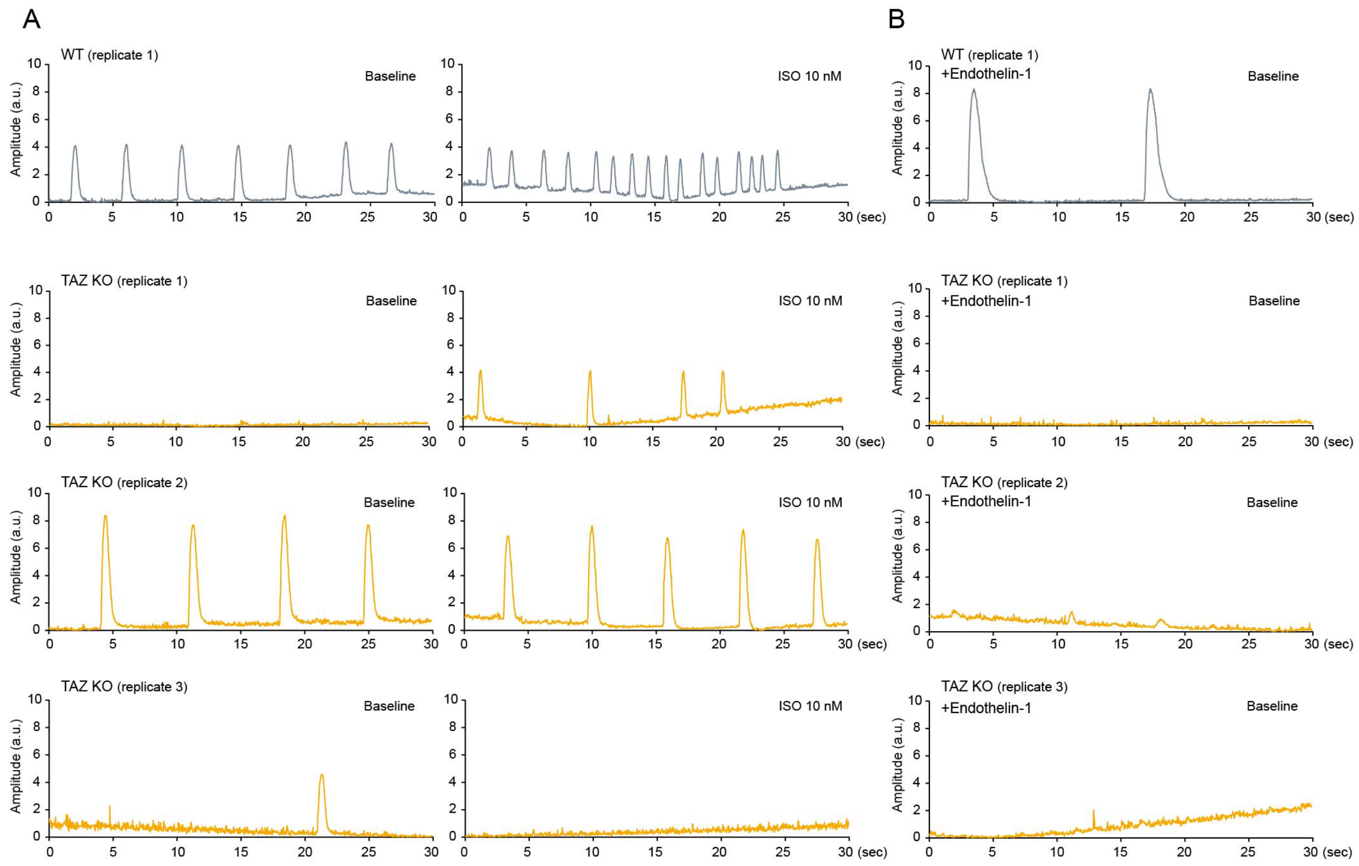


**Figure S13: Contraction profiles of WT and TAZ KO iPSC-derived cardiomyocytes.**

Related to Figure 6, representative traces of cardiomyocyte contractions over time for WT and TAZ KO cardiomyocytes cultured with Pal/car + Dex/T3 for 2 weeks, in the absence (A) or presence (B) of 10 nM endothelin-1.

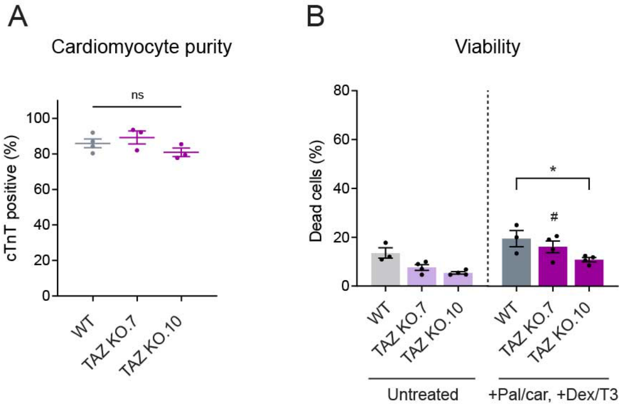


**Figure S14: Purity and viability of C11 WT and TAZ KO iPSC-derived cardiomyocytes.**

**(A)** Cardiomyocyte purity, presented as cardiac troponin T (cTnT) positive population. C11 WT and TAZ KO cardiomyocytes at baseline were analyzed (mean ± SEM, n = 3-4 biological replicates). **(B)** Viability of C11 WT and TAZ KO cardiomyocytes after 2 weeks of treatment with Pal/car + Dex/T3 (mean ± SEM, n = 3-4 biological replicates). Statistical significance: one-way ANOVA with Tukey's test; *$p<0.05$. * WT vs. TAZ KO within treatment; # vs. untreated within genotype.

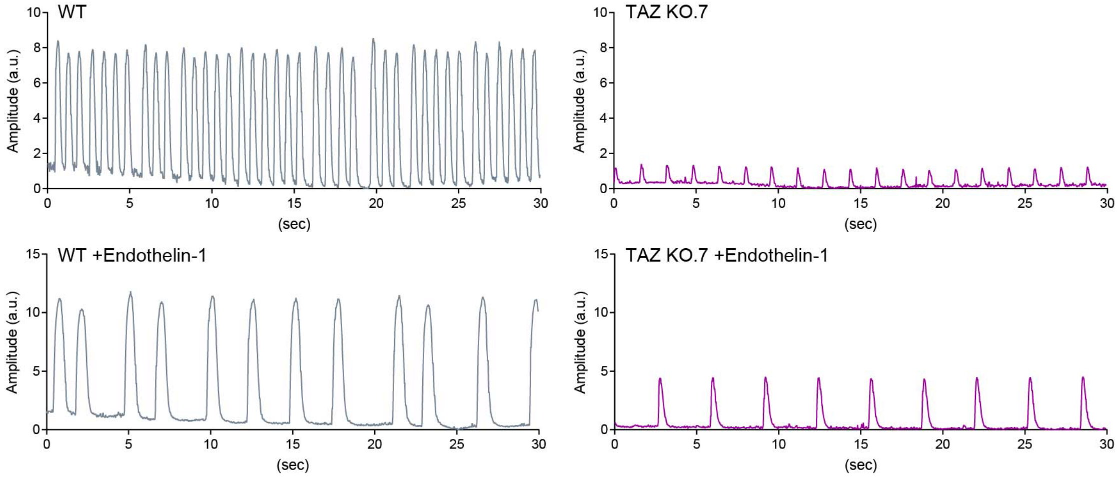


**Figure S15: Contraction profiles of C11 WT and TAZ KO iPSC-derived cardiomyocytes.**

Related to Figure 7, representative traces of cardiomyocyte contractions at baseline for C11 WT and TAZ KO cardiomyocytes cultured with Pal/car + Dex/T3 for 2 weeks, in the absence or presence of 10 nM endothelin-1.

**Table S1: qPCR primers used in this study**

| Gene | Forward (5'-3') | Reverse (5'-3') |
|---|---|---|
| YWHAZ | TGCTTGCATCCCACAGACTA | AGGCAGACAATGACAGACCA |
| EEF1A1 | CTGGACTGCATCCTACCACC | CTCGGCCAACAGGAACAGTA |
| MYL2 | TGTCCCTACCTTGTCTGTTAGCCA | ATTGGAACATGGCCTCTGGATGGA |
| TNNI3 | CGTGTGGACAAGGTGGATGA | CCGCTTAAACTTGCCTCGAA |
| TNNI1 | TGGTGGATGAGGAGCGATAC | GTCCATCACCTTCAGCTTCAG |
| CACNA1C | AAGGCTACCTGGATTGGATCAC | GCCACGTTTTCGGTGTTGAC |
| ATP2A2 | CAGCCTTTGTAGAACCTTTTGT | AATCCGCTGCACACTCTTTC |
| KCNJ2 | TGTCACGGATGAATGCCCAA | CAAACACAGCTTGCCGTCTC |
| NPPA | CCGTGAGCTTCCTCCTTTTA | CCAAATGGTCCAGCAAATTC |
| NPPB | CTCCAGAGACATGGATCCCC | GTTGCGCTGCTCCTGTAAC |
| PRELID | ACCAAGTGTTCGCCGCCTTCTG | GCCGGGACAGCAGTTTCTGGTC |
| TRIAP1 | CCCGTGCACCGACCTCTTCAAG | TGCCATGGCCCATGAACTCCAG |
| TAMM41 | GTCTACGGCTCCGGGGTGTAC | GCCATGCGACAGGGTCATCTAC |
| PGS1 | GCCAAGAGGCGGGTCGTGATG | TCGTGAGCCCCGCGTGAAG |
| PTPMT1 | CTGCCGTTGCGGAGCTTGA | CGCAGCTGCTCGACTCCTAGT |
| CRLS1 | ACACCACGAACACTTGCCAAGT | TGATGCAGCTGTGGTGAAAGCT |
| TAZ | TACATGAACCACCTGACCGT | CAGATGTGGCGGAGTTTCAG |
| PNPLA8 | TGCTGCTCCAGGCTACTTTGCA | AACGTCCAGTGCCCAGGGATAC |
| LCLAT1 | CAACCGCCTTGTGGCAACATG | TCGCATCAGGCAATTCCACAGG |
| ABHD18 | GAGGATGGGGAAGGCCAGAAGA | TGAGCCATGGGGGAAACAAAGTG |
| HADHA | CCTGCCTGGGAGGAGGACTTG | CAGCAGGCACACCCACCATT |
| MT-ND1 | ACGGGCTACTACAACCCTTC | GCCTAGGTTGACCA |
| B2M | GGAGAGCTGTGGACTTCGTC | ATTCTACAAACGTCGCGTGC |
